# Optical recordings of synaptic vesicle fusion reveal diffusional dispersion

**DOI:** 10.1101/2025.03.31.646326

**Authors:** Julia Lehrich, Nataliya Glyvuk, Junxiu Duan, Sai Krishnan, Yaroslav Tsytsyura, Ulrike Keller, Lakshmi Penothil Sunil, Georgii Nosov, Jana Hüve, Philipp Selenschik, Carsten Reißner, Markus Missler, Martin Kahms, Jürgen Klingauf

## Abstract

The primary unit of exocytosis, the synaptic vesicle, is replete with proteins which enable vesicle fusion. For maintaining synaptic transmission the vesicle proteins have to be rapidly cleared from the active zone. The immediate fate of the cargo post-fusion and the precise mechanism of its resorting into a readily retrievable pool (RRetP) for endocytosis remains unclear. To address this, we developed a purely presynaptic preparation, the xenapse, presynaptic boutons formed *en face* directly onto a glass coverslip enabling optical single vesicle recording by total internal reflection microscopy (TIRFM). Electron microscopy showed xenapses contain a few hundred synaptic vesicles (SVs), of which ∼40 SVs are primed (readily releasable pool, RRP), as revealed by single vesicle recordings using the pH-sensitive fluorescent protein pHluorin. Single fusion events synchronous with action potentials could be localized with ∼20 nm precision and rapid post-fusion diffusional dispersion of vesicular proteins observed with diffusion constants in the order of 0.1 µm^2^/s. Unroofing of xenapses revealed numerous Clathrin-coated structures, enriched for vesicle cargo and numerically equivalent to about twice the measured RRP size. In this way SV proteins are rapidly cleared from the release site into this RRetP of pre-assembled pits to allow for rapid re-docking and re-priming at vacated release sites in synapses tuned for high-fidelity fast signaling.

## Main

Different mechanisms have been proposed for SV fusion and for the subsequent retrieval of SV proteins, the cargo. Vesicles might release their content by connecting transiently to the plasma membrane without full collapse and loss of protein content (‘kiss-and-run’)^1–5^. While this maintains the vesicle integrity, the major release pathway involves the full-collapse fusion of the vesicle onto the membrane. The neuron then faces the challenge of clearing the SV components from the release site and fully reconstituting them into new fusion-competent vesicles^6–10^. It can do this by Clathrin-mediated reclustering, resorting and endocytosis of vesicle proteins ^11–14^. Alternatively, after full vesicle collapse, SV proteins and lipids do not disperse in the plasma membrane but remain clustered by self-assembly in raft-like structures which could be endocytosed *in toto* ^15^. The relative contribution of these different mechanisms to compensatory endocytosis in the CNS however remains ambiguous. Recent results obtained by electron microscopy after rapid high-pressure freezing suggested the existence of an ultrafast endocytosis (UFE) mechanism, at a spatially distinct periactive zone, starting 50–100 ms after stimulation ^16,17^, where SV membrane is retrieved into large vesicles of four-fold membrane area compared to SVs. It is conceivable that such a rapid retrieval might favor raft-like structures of SV proteins that remain clustered by self-assembly after SVs collapse and flatten within the PM, although a sub-second retrieval mechanism in frog neuromuscular junction involving large vesicles of the same size as UFE has been shown to not sort cargo, and was attributed to unphysiological stimulation ^14^. On the other hand, maintaining molecular identity during exo-endocytic cycling is the hallmark of a kiss-and-run-type mechanism. The fate of SV proteins post-fusion is thus intimately connected to the mechanism of endocytosis the neuron might use to retrieve vesicles.

While ‘kiss-and-run’ does not leave behind a cargo footprint on the membrane, both modes of full collapse can be easily distinguished by the dispersion criterion at the single vesicle level. The loss of molecular identity during exo-endocytosis triggered by short stimulus trains was shown ^9,18,19^ using fusion constructs of the pH-sensitive GFP variant pHluorin (pHl) ^20^ with the vesicle SNARE synaptobrevin-2 (SynaptopHluorin, Syb2-pHl) or the putative Ca^2+^ sensor Synaptotagmin-1 (Syt1-pHl) ^9^. We showed in previous studies that a large fraction of these surface-stranded SV proteins are pre-sorted and pre-assembled in a so-called readily retrievable pool (RRetP) of synaptic vesicle proteins ^9,10^.

Despite these advances in our understanding of cargo clearance, resolving the fine details of this process has eluded us due to the inability to visualize and track the fate of individual cargo. The primary reason for this limitation arises from the inaccessibility of small CNS boutons, compounded by the random orientation of the AZ in *en-passant* synapses in neuronal cultures. The earliest attempts of overcoming this problem used freshly isolated terminals of goldfish retinal bipolar cells, calyces of Held, as well as hippocampal mossy fiber synapses ^21–23^. The terminals of these cell preparations can be placed face down on coverslips and TIRFM used to enhance contrast. All these acutely isolated cell preparations, however, survive only a few hours and thus cannot be labelled genetically, e.g. by pHl constructs.

More recently, various attempts have been made to enrich for presynaptic structures by immobilizing post-synaptic synaptic cell adhesion molecules (sCAMs) like Neuroligin-1 (Nlgn1) or synCAM ^24,25^ onto a solid surface ^26–28^. But achieving TIRF-amenability rests on precisely defining the formation of the synaptic structure in two dimensions directly on the substrate. While axonal swellings have been generated to display “active zone-like” features, no single AP-secretion coupling was observed. This lack is a sign of immaturity, likely arising from synaptogenic sCAMs being immobilized on the surface using antibodies or biotin-streptavidin, which are bulky and limit the density of protein labelling. The real stumbling block to inducing robust TIRF-amenable presynapses therefore has been the failure so far to immobilize the synaptogenic protein with sufficiently high density.

We sought to overcome this bottleneck by developing a novel antibody-free and biotin-streptavidin-free approach of micropatterning Nlgn1 onto a glass coverslip. This resulted in the ‘xenapse’, presynaptic boutons formed by cultured mouse hippocampal neurons on functionalized micropatterned host (old greek: xenos) substrates. Xenapses show the hallmarks of a typical mature pre-synapse both structurally (SV clusters, distinct exocytic and endocytic zones) and physiologically (synchronous and ‘phasic’ release). We show that proteins disperse rapidly by diffusion after exocytosis, disproving the long-standing controversial ‘kiss-and-run’ hypothesis. Measurements at unprecedented detail revealing the spreading of several SV molecules were only made possible by the xenapses being sufficiently robust for single vesicle recordings by TIRFM. Regarding assembly of the RRetP, we found that self-assembly of SV proteins is not sufficient, but that Clathrin, adaptor proteins and other accessory proteins are additionally required. Platinum replicas of cell cortices produced by ‘unroofing’ of xenapses revealed numerous pre-assembled Clathrin-coated structures representing the RRetP and counterbalancing the RRP.

### Xenapses, presynaptic hippocampal boutons on micropatterned coverslips

Nlgn1, which binds the presynaptic sCAM Neurexin (Nrxn), has been demonstrated to be sufficient for inducing formation of presynapses even when overexpressed in non-neuronal cells ^24,29–31 25,30–32^, with Liprin-α proteins linking *trans*-synaptic Nlgn1-Nrxn contacts to downstream steps ^31,32^. We were able to induce hippocampal neurons to form purely presynaptic structures ‘*en face*’ directly onto the coverslip at predefined spots in a grid-like pattern (Fig. 1a). Grid patterns (5 µm diameter, 8 µm center to center distance, 55 % coverage) of a polymer nano-brush using a polyethylene glycol (PEG) backbone for the brush are formed by micro-contact printing (µCP) with polydimethylsiloxane (PDMS) stamps ^33^ (Fig. 1b). For inducing synaptic contacts a pre-synthesized poly-l-lysin-PEG (PLL-PEG) brush polymer grafted with HaloTag ligand (HTL) was used (grafting ratio 3.5, height of about 1 – 2.5 nm, Fig. 1c), while a PLL-PEG-methoxy brush was used for backfilling. Incubation with a purified fusion protein of the extracellular Nlgn1 domain (splice variant +A-B) and the Halo protein led to strong functionalization of grid points as evaluated by immuno-fluorescence staining (Fig. 1d). Formation and direct interfacing of these purely presynaptic boutons on the micropatterned host substrate (a polymer nano-brush: PLL-PEG-HTL, cf. Methods), which we call ‘xenapses’, are induced shortly (DIV 1-2) after cell seeding (suppl. Fig. 1a). No xenapse formation was observed when grafting the RGD domain of the ubiquitous cell adhesion protein Integrin to the nano-brush (data not shown). Xenapses can be scaled in size simply by varying grid point size, (e.g. spot size 2.5 µm, 3 µm center to center, coverage 23 %, Suppl. Fig. 1e). Empirically, we found, however, that µCP becomes unreliable with stamp spot sizes smaller than 1.5 um, because of insufficient stiffness of the PDMS stamp brush. Xenapses matured within 2 weeks (Fig. 1e), and were considered mature when forming AZs (cf. Fig. 3) within two weeks, and responding to single elicited APs with synchronous release of SVs (cf. Fig. 4). No mature xenapse formation, using the above criteria, was observed when using a striped pattern instead of a grid pattern, irrespective of the coverage (suppl. Fig. 1b,c,d), only elongated swellings were formed by axons on the functionalized stripes, that barely responded to stimulation. While dendrites meander rather randomly on the methoxy- and PLL-PEG-HTL substrate and form conventional synapses with crossing axons (Fig. 1f), many axons directly connect functionalized grid points and form large boutons *en face* (Fig.1g). Both, glutamatergic and GABAergic boutons are formed at a ratio of about 2:3 (suppl. Fig.1f,g), although the Nlgn1 splice variant used is the one with the highest affinity for Neurexins (Nrxn) and is most prevalent in inhibitory neurons ^34^.

**Figure 1.**
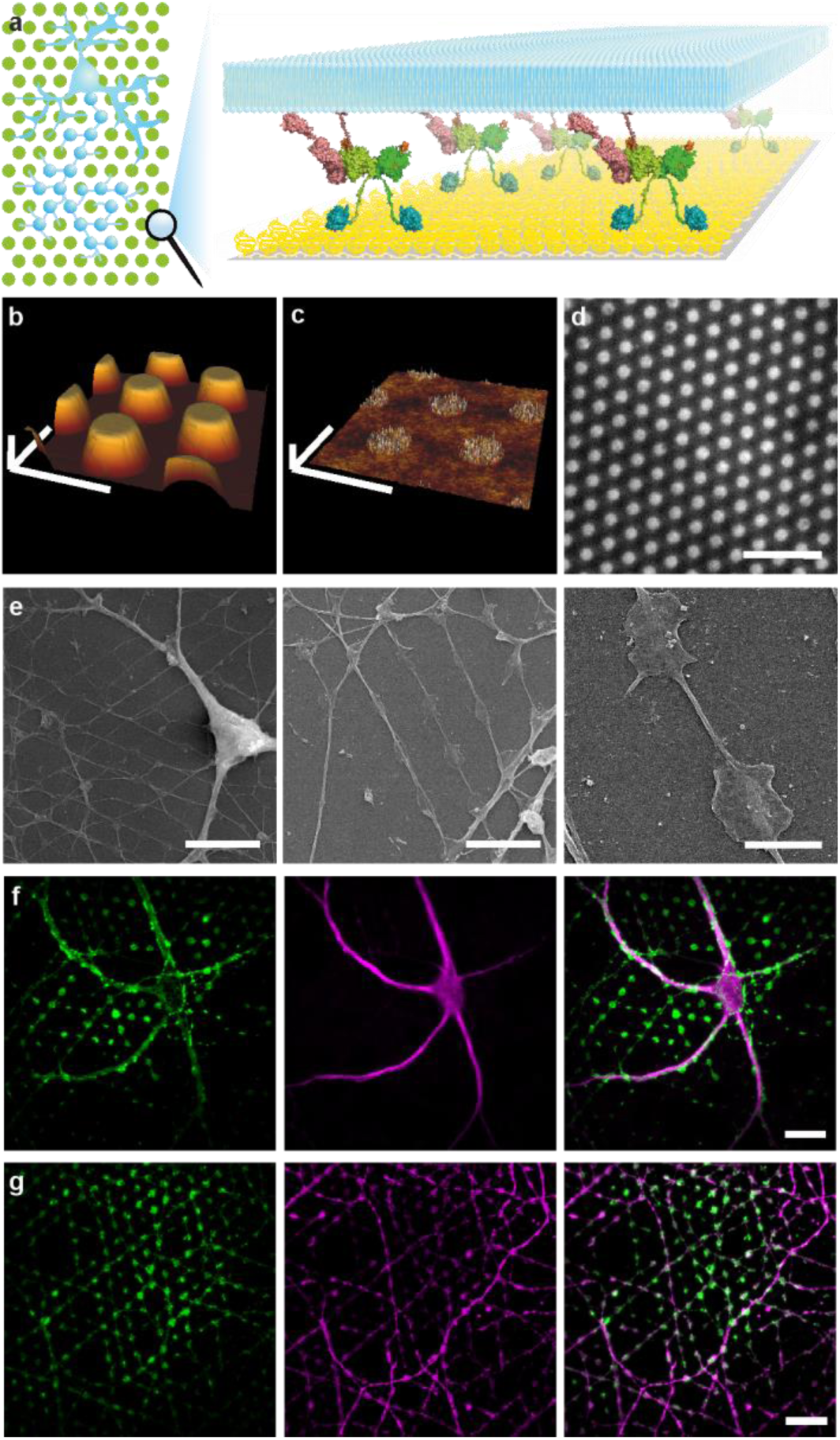
Microstructured coverslips functionalized with the synaptic cell adhesion molecule Nlgn1 serve as artificial host substrates to induce presynapses. **a**, Left, cartoon of a neuron growing on a substrate functionalized with Nlgn1 (green dots) and, right, schematic of the patterned substrate (PLL-PEG-HTL, yellow) with presynaptic membrane of a xenapse (plasma membrane, light blue) attached via Nrxn1-Nlgn1 complex (quaternary structure, pink and green) and the HaloTag linker (dark blue). **b and c**, AFM images of the topology of a polymer stamp used for microstructuring (**b**) and the microstructured substrate (**c**). Scale bars 8 µm (x and y) and **b**, 800 nm (z) or **c**, 10 nm (z). **d**, Surface functionalized Nlgn1 visualized by antibody staining, scale bar 25 μm. **e**, Scanning electron microscopy images of hippocampal neurons (9 DIV) growing on the functionalized substrate with zoom-in of a single axon forming two xenapses (right). Scale bars 20 µm, 12 µm and 3 µm, respectively. **f and g**, Confocal image of xenapses stained for vesicle protein Syp1 (green) and the dendritic protein MAP2 (magenta, **f**) or the axonal protein tau (magenta, **g**) indicating varicosities on the functionalized substrate originating from axons only. Scale bars 20 μm.

### Xenapses show structural hallmarks of conventional synapses

Horizontal sections by FIB-SEM (Fig. 2a, suppl. Fig. 2) display numerous physically docked SVs and show that the synaptic cleft between the polymer brush nanostructures and xenapses is ∼ 15 nm. This is the same size as in the standard bouton counterparts, as cleft width is dictated by the transsynaptic synaptic cell-adhesion complexes, in our case the Nlgn-Nrxn complex. 3D reconstructions of electron micrographs of serial sections of large xenapses reveal multiple clusters of docked SVs and show that xenapses harbor several mitochondria (Fig. 2b). Comparison to normal cultured hippocampal boutons (Fig. 2c) reveals similar numbers of SVs visibly docked to the plasma membrane per AZ membrane area (Fig. 2d), comparable densities of SVs in the clusters above the AZ (Fig. 2e), and identical SV sizes (Fig. 2f).

**Figure 2.**
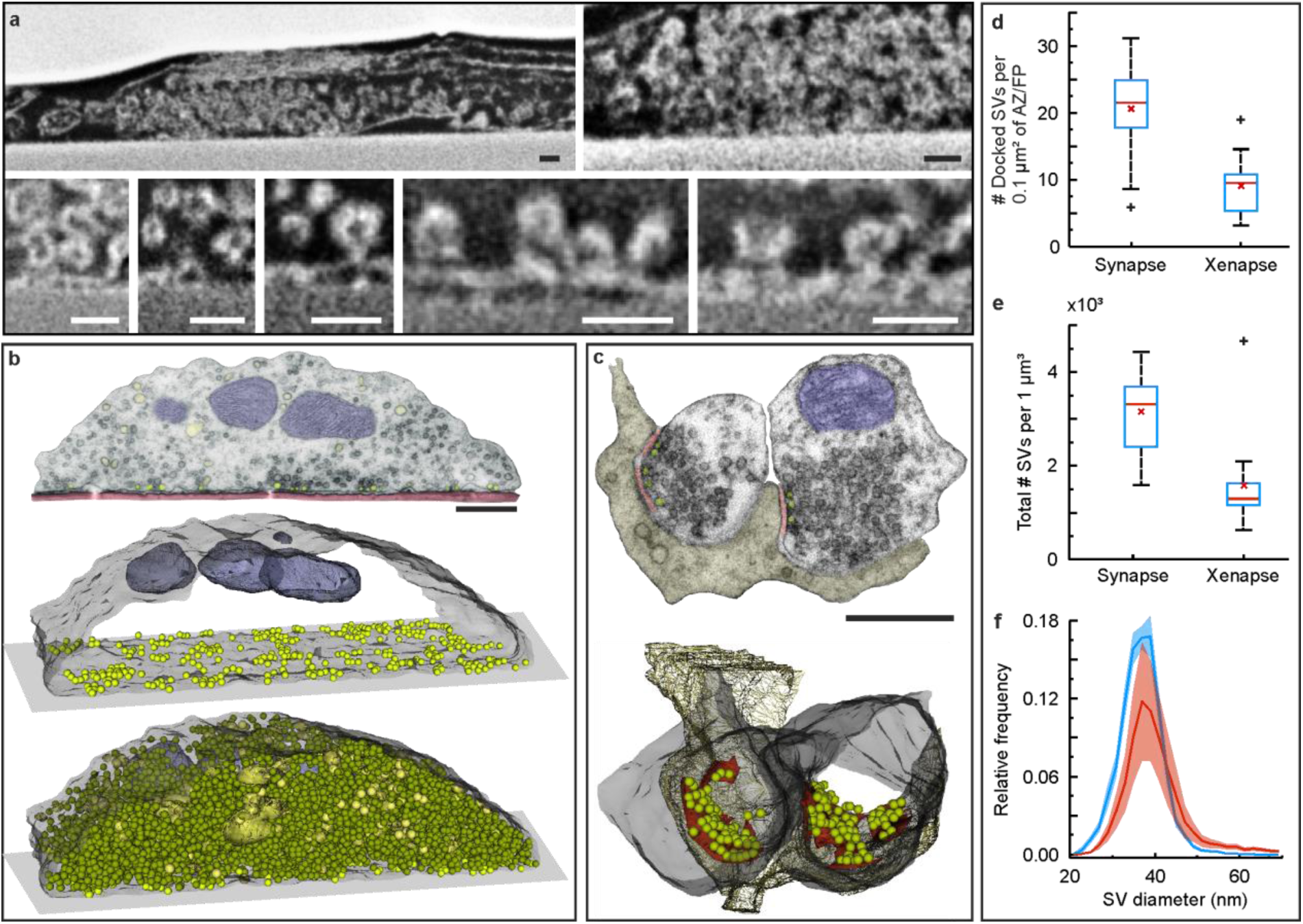
Xenapses form directly on the functionalized substrate with many vesicles docked at the bottom facing the Nlgn1 protein pattern. **a**, FIB-SEM sections of xenapses show the intact cell-substrate interface. SVs can be found in areas where xenapses are tightly attached to the substrate. Scale bars 100 nm. **b**, 3D reconstruction of TEM serial sections retrieved from a single xenapse (16 DIV); top panel: representative serial TEM image. Scale bar 500 nm; middle panel: the contour of the xenapse (light gray) with docked synaptic vesicles (light green) at the plasma membrane contacting the glass coverslip (flat rectangle at the bottom of the xenapse) and mitochondria (violet-grey) are displayed; bottom panel: total pool of synaptic vesicles (green). **c**, 3D reconstruction of TEM serial sections of normal cultured hippocampal boutons. Color coding as in **b**. Scale bar 500 nm. **d**, Number of docked SVs per active zone area of the synapse (n=26) or per footprint area of the xenapse (n = 13). **e**, Number of SVs per volume of synapse (n = 18) or xenapse (n = 13). **f**, Synaptic vesicle size distribution for synaptic (blue, n = 7075 SVs, N = 12 synapses) and xenaptic (red, n = 18619 SVs, N = 4 xenapses) boutons.

### Xenapses harbor about 16 AZs of similar size as in synapses

TIRF-dSTORM superresolution images of xenapses stained for the abundant SV protein Synaptophysin-1 (Syp1) to label SV clusters and for the large scaffold protein Bassoon as a marker for AZs display several hotspots (Fig. 3a). A good marker of AZs and hotspots of docked and primed SVs is the SV priming factor Munc13-1 ^35,36^. Thus, we used a CRISPR/Cas9-based knock-in approach to tag endogenous Munc13 with EGFP ^37^. Fluorescent signals were very faint and thus enhanced by nanobody-staining. Confocal and STED imaging revealed colocalizaion with the bright Bassoon clusters (Fig. 3b,c). We next analyzed reconstructed STORM superresolution images of Bassoon localizations by Voronoi tessellation ^38^ (Fig. 3d), which yielded a median value of 16 AZs in xenapses growing on 5 µm spots with a mean area corresponding to a circle of 199 nm diameter (Fig. 3f,g), similar to AZ size in hippocampal synapses ^39^. Xenapses grown on 2.5 µm spots harbor a median of 7 AZs of same size, suggesting that AZ size is rather intrinsically regulated and less dependent on the Nlgn1-functionalized spot size (Fig. 3e,f,g).

**Figure 3.**
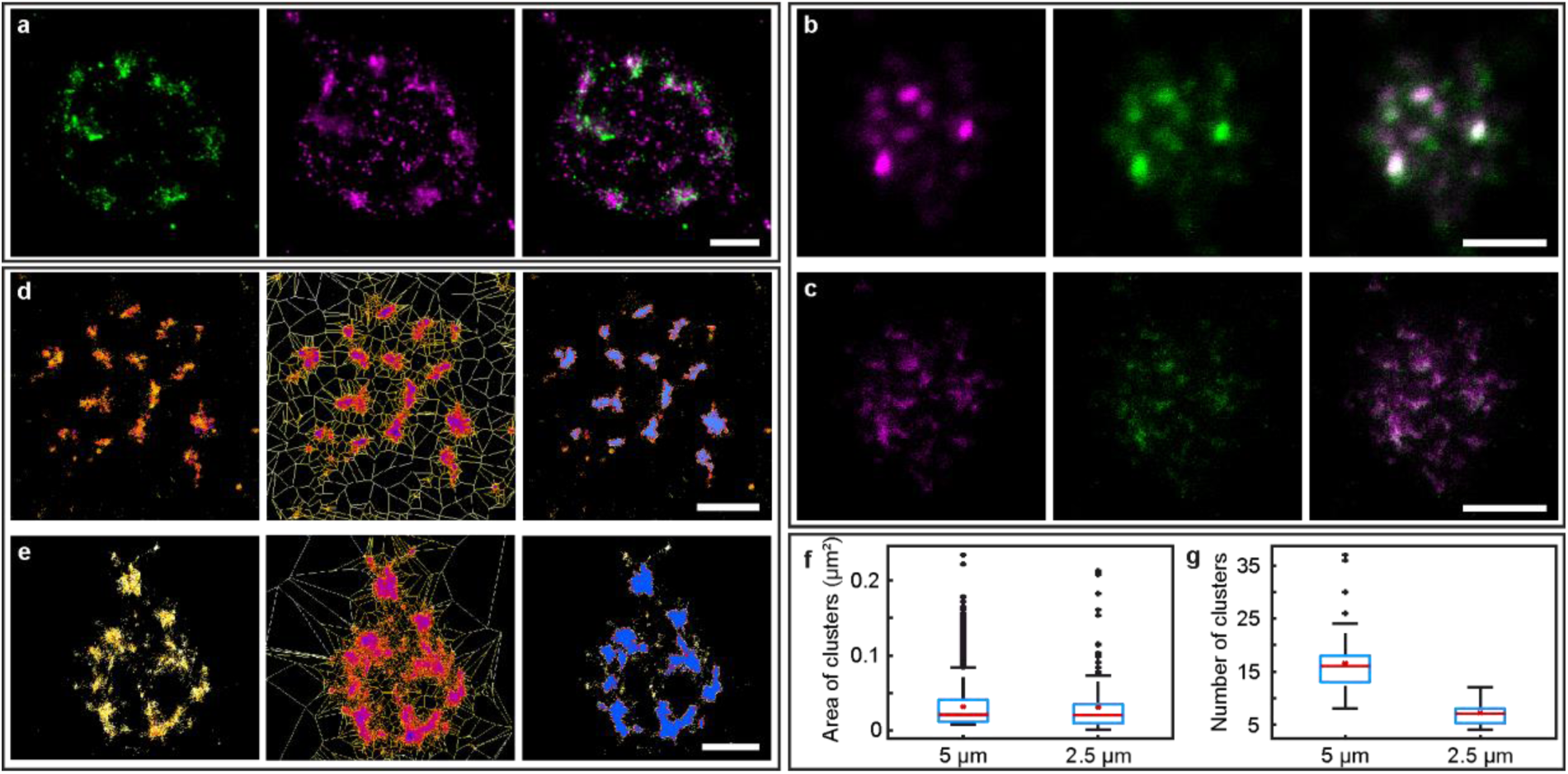
Xenapses form many AZs. **a**, TIRF-dSTORM images of an exemplary single xenapse with Syp1 (green, left) and Bassoon (magenta, center, and merge, right) labelling. Xenapses feature several protein clusters, indicating the presence of multiple active zones. Scale bar 1 µm. **b and c**, Xenapses growing on 5 µm spots expressing Munc13-GFP KI imaged using, **b**, confocal and, **c**, STED images (green: Munc13 KI/Anti-GFP nb 635P, magenta: Bassoon) showing colocalization of Munc13 and Bassoon. Pearson correlation coefficient was determined for confocal (R = 0.8184, 88 xenapses) and STED images (R = 0.5841, 45 xenapses). Scale bar 2 µm. **d and e**, Bassoon cluster analysis of reconstructed STORM image (left), Voronoi tessellation (center) and identified clusters for xenapses on 5 µm (**d**) and 2.5 µm spots (**e**). Scale bar 1 µm. **f**, Distribution of the area of Bassoon clusters for xenapses growing on 5 µm spots (n = 1870 clusters from 113 xenapses in 7 regions from 4 biological replicates) and on 2.5 µm spots (n = 221 clusters from 31 xenapses in 3 regions from 3 biological replicates). **g**, Distribution of the number of Bassoon clusters for xenapses growing on 5 µm spots (mean of 16.5, n = 113 xenapses, 7 regions and 3 biological replicates) and on 2.5 µm spots (mean of 7.2, n = 31 xenapes, 3 regions and 3 biological replicates).

### Xenapses display exo-endocytosis, stimulus train responses and RRP sizes like synapses

Next, we investigated the physiological properties of xenapses. We loaded xenapses with the exogenous low affinity fluorescent Ca^2+^ sensor Fluo-5F and elicited single APs at 2 Hz by short 1 ms electric field pulses in TIRF illumination. As in their small synaptic counterparts robust Ca^2+^ influx could be observed for every AP (Fig. 4a,b). To assess exo-endocytosis we expressed Syp1 harboring two pHl (Syp1-2xpHl) in xenapses. When trains of 50 and 100 APs at 20 Hz were applied, exocytosis of Syp1-2xpHl in synchrony with stimulation was observed, followed by fully compensatory endocytosis with time constants of 15.1 s and 27.5 s at room temperature (Fig. 4c,d). When APs are spaced by at least 3 frames, i.e. 30 ms, individual fusion events can be resolved in space and time (Fig. 4e,f). Two different responses of xenapses can be observed under these conditions. While some xenapses showed initial facilitation for the second stimuli (Fig. 4g), followed by depression down to a steady-state level of sustained release, most xenapses displayed immediate depression (suppl. Fig.4c,d), often with a very high release response to the first AP, reminiscent of other ‘phasic’ synaptic preparations ^40^. The same behavior was observed in small hippocampal synapses expressing Syp-2xpHl (suppl. Fig. 4a,b). Xenaptic responses could be increased by application of the DAG analog Phorbol 12, 13-Dibutyrate (PDBu) (suppl. Fig. 4e), which activates Munc13 and the kinase PKC ^35^ and thereby increases the number of release-ready SVs like in other CNS synapses. We next attempted to quantify the RRP size in xenapses. To this end, we applied trains of 40 APs at 33.3 Hz (Fig. 4e,f,g). At higher frequencies, short-term depression (STD), likely due to limited release site clearance, is observed ^41^, which would confound RRP size estimates. From such measurements, the total RRP size of xenapses can be estimated from plots of cumulative release against stimulus number by back-extrapolation of a regression line to the late sustained phase, assuming a constant pool replenishment ^42^. This method requires conversion of the release amplitudes into numbers of released SVs. While this is impossible by pHl measurements in small hippocampal boutons due to the lack of spatial resolution and due to the broad distribution of pHl molecules per SV ^43^, individual fusion events can be spatially well resolved in TIRFM recordings of large xenapses, at least for the first stimulus, when there is no major buildup of exocytosed pHl on the surface yet (Fig. 4f). When calibrating release amplitudes in this way (Fig. 4h), we find for all xenapses, i.e. facilitating and depressing ones, an average RRP size of about 40 SVs using the back-extrapolation method (Fig. 4i). With on average 16 AZs per xenapse (Fig. 3) this amounts to 2.5 release sites per AZ, close to the estimate of about 3 for small hippocampal boutons in culture ^44^, the calyx of Held ^45^ and a CNS synapse ^46^.

**Figure 4.**
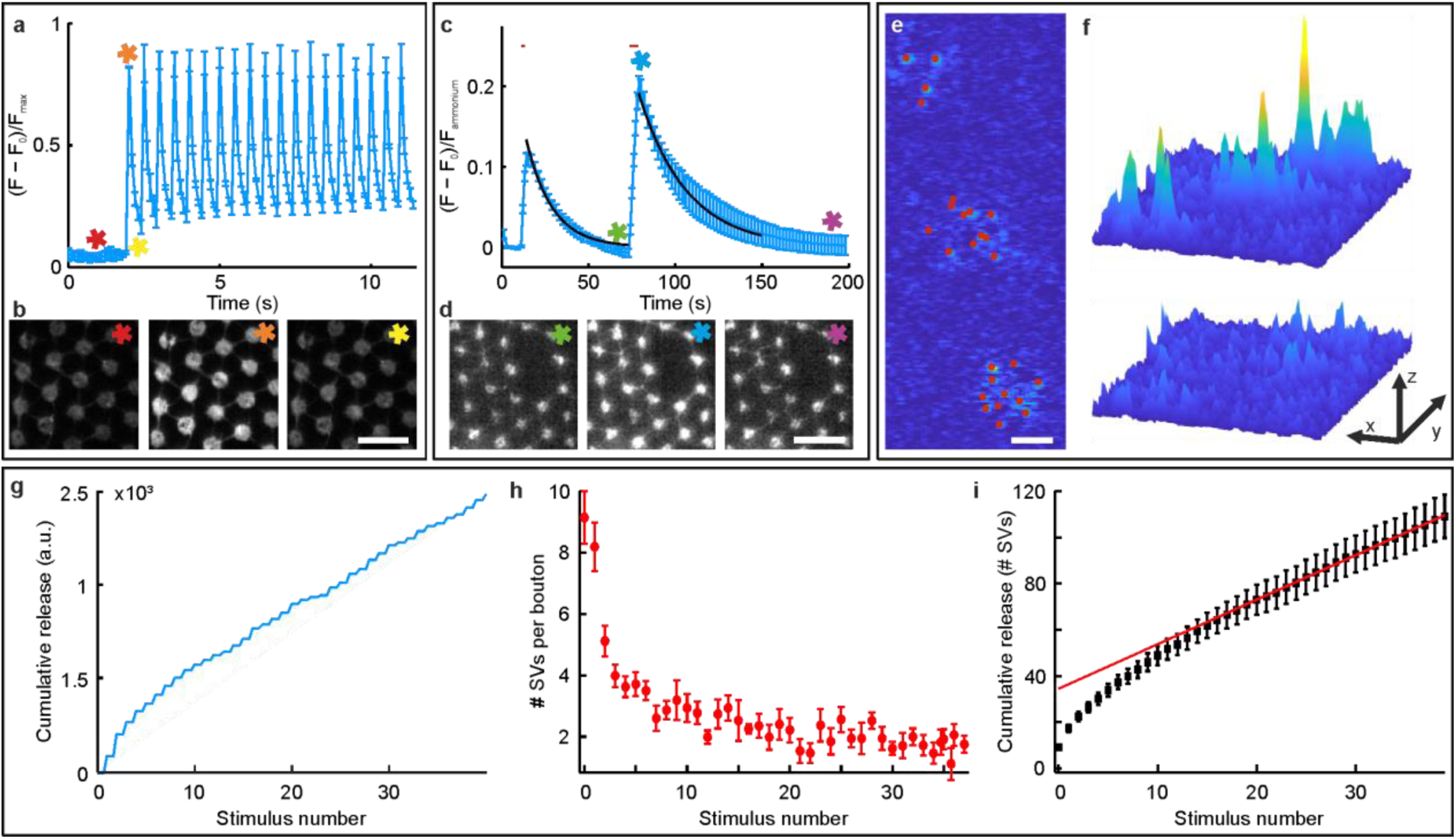
Xenapses display Ca^2+^ entry, exo-endocytosis, and RRP sizes similar to synapses. **a**, Average fluorescence signal of xenapses (DIV 15) loaded with the Ca^2+^ indicator Fluo5F in response to single APs elicited at 2 Hz. Error bars in s.e.m.. **b**, Single representative images captured at color-coded time points: asterisks mark images before (red), during (orange) and after (yellow) stimulation (15 DIV, n=6 regions with tens of xenapses each, 3 biological replicates). Scale bar 12 µm. **c**, Syp1-2xpHl recordings of xenapses stimulated with 50 and 100 APs at 20 Hz with fluorescence intensity profiles normalized to maximum intensities during NH_4_Cl superfusion after stimulation. (n = 6 regions with tens of xenapses each, 3 biological replicates). Error bars in s.e.m., red bars indicate stimulation pulses. **d**, Single images taken at color-coded time points: asterisks mark images before (green), during (blue) and after (violet) the second pulse of stimulation. Scale bar 12 µm. **e**, TIRFM image of single SV fusion events detected with Syp1-2xpHl upon a single AP in three adjacent xenapses. Single events (red circles) were fit by Gaussians and the fits were subtracted from the first image after the action potential in an iterative process to also detect partially overlapping events. Scale bar 2 μm. **f**, Surface plot of the SV fusion events in **e**. After three iterations no further events could be detected in the subtracted image (bottom). Scale bars 2 μm (x), 11 μm (y) and 100 a.u. (z). **g**, Cumulative responses to a train of 40 APs at 33 Hz of all 21 xenapses from the same measurement. **h**, Average responses to 40 APs (n = 6 regions each comprising > 15 xenapses, N = 4 biological replicates). Using the detection algorithm in (**e**) numbers of released SVs were determined for the response to the first AP and used to convert fluorescence increases upon all subsequent APs into numbers of released SVs per xenapse. Errors in s.e.m.. **i**, Cumulative plot of responses in (**h**) and regression fit to the last 20 responses for determining the average RRP size per xenapse.

### Optical recordings of single fusing SVs allows quantifying the pHl content per SV

In TIRF illumination, xenapses expressing Syp1-2xpHl were stimulated by a few 10 APs at 2 Hz. Prior to stimulation most of the orphan surface-stranded pHl molecules were prebleached to enhance the contrast and signal to noise ratio of single fusion events. Images show spatially resolved individual fusion events upon single APs (suppl. Fig. 4f), which could be localized using template-based Gaussian fits with a precision of about 20 nm. Calibration of the scientific Complementary Metal-Oxide-Semiconductor (sCMOS) chip (cf. Methods) allowed us to quantify the number of detected photons for each individual exocytosis event (Fig. 5f). An amplitude histogram of 1188 individual events yields a skewed distribution with equidistant subpeaks or shoulders (suppl. Fig. 4g), as we also have previously observed in small synaptic boutons ^43^, raising the question, whether these subpeaks and shoulders correspond to individual pHl molecules. We thus immobilized purified EGFP as well as pHl to coverslips and measured individual bleaching steps on the same TIRFM setup. Photon histograms of amplitudes could be fitted with single Gaussians, yielding average amplitudes of 359.76 ± 139.34 photons for EGFP and 280.22 ± 129.89 photons for pHl (suppl. Fig. 4h). Interestingly, spatially resolved blinking events prior to stimulation, i.e. during baseline recording, showed a near-identical Gaussian distribution with an average of 303.1 ± 86.2 photons (suppl. Fig. 4g, blue). Thus, these events probably arise from single blinking of surface-stranded pHl molecules out of a dark state under these rather high laser power conditions (196 mW/cm^2^ at the focal plane). Recalibration of the release event amplitude histogram to numbers of pHl molecules using the average photon count (suppl. Fig. 4g, upper abscissa) indeed confirms, that equidistant subpeaks and shoulders arise from the contributions of individual pHl molecules. Since Syp-2xpHl, however, contains two pHl moieties, we repeated these experiments using xenapses expressing Syt1-pHl, which contains only one pHl moiety (suppl. Fig. 4i). This histogram confirms earlier calibrations of single SV pHl responses in small hippocampal boutons, which revealed that most SVs harbor only 1 to 2 pHl molecules ^43,47^. This low pHl copy number, however, hampers the analysis of the fate of SV constituents post fusion. Since xenapses allow the recording of hundreds to thousands of spatially resolved single release events, one can restrict the analysis to those SVs that at least contain two pHl-tagged proteins, i.e. four pHl molecules for Syp1-2xpHl and two for Syt1-pHl (suppl. Fig. 4g,i, dashed blue lines).

**Figure 5.**
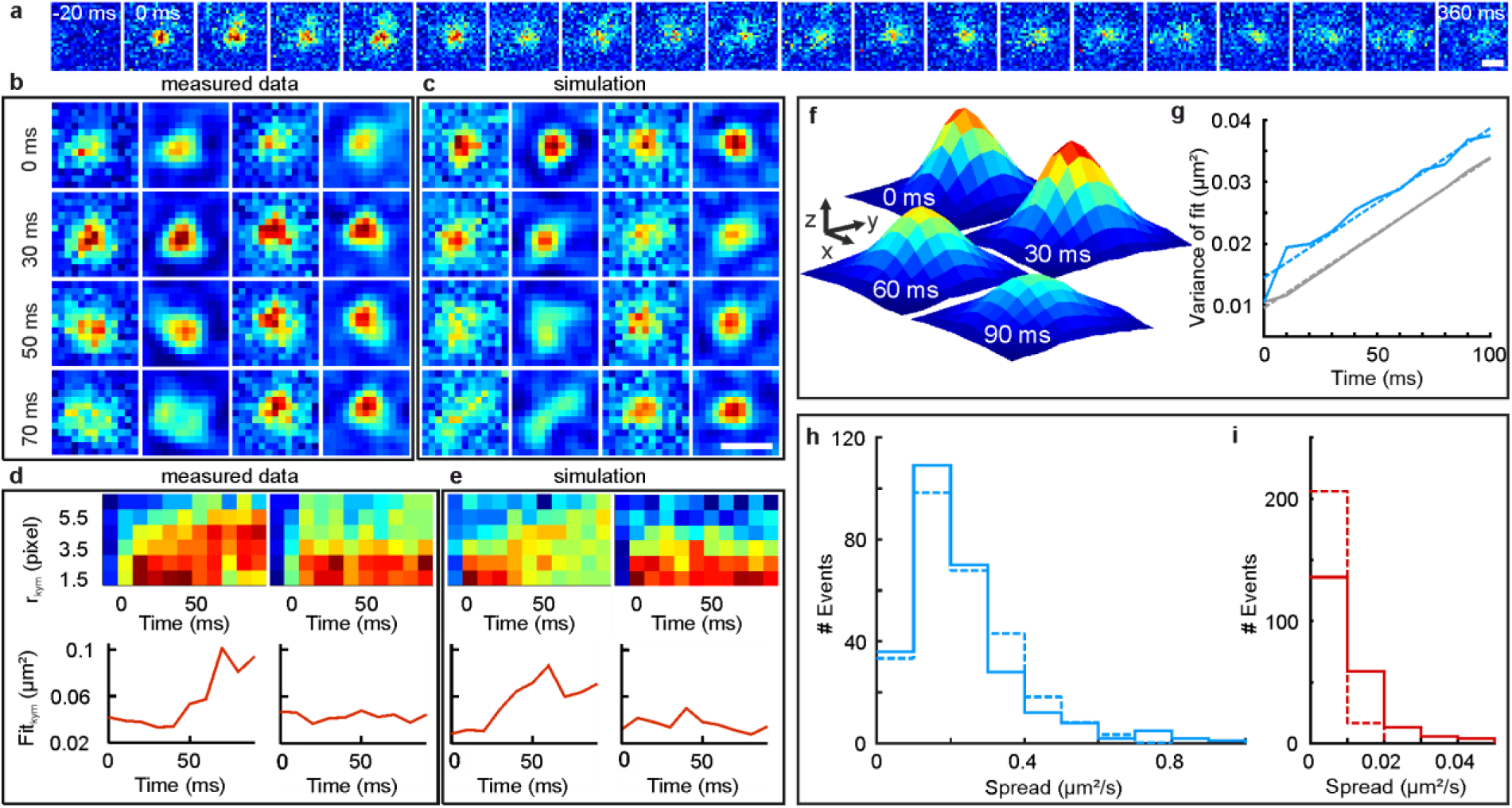
Recordings of single SVs harboring multiple Syp1-2xpHl reveal diffusional dispersion of SV proteins post fusion. **a**, The fluorescence spread of single fusion event recorded and tracked over 360 ms. Scale bar 500 nm. **b**, Two exemplary fusion events, with raw data and wavelet filtered data. Left: Fusion event with wide spread of fluorescence post-fusion (fast diffusion). Right: fusion event with narrow spread (slow diffusion). **c**, Simulated fusion events (raw data and wavelet filtered data) with wide spread of fluorescence post-fusion (fast diffusion, left), and narrow spread (slow diffusion, right). Scale bar 500 nm. **d**, Quantification of the spread of rapidly dispersing (upper left) or slowly dispersing measured fusion events (upper right). The time-dependent change of the width at half maximum is displayed for rapidly spreading (lower left), or slowly spreading fusion events (lower right). **e**, Radial kymographs of rapidly spreading (upper left), or slowly spreading simulated events (upper right), as well as the time-dependent change of the width at half maximum for rapidly spreading (lower left) or slowly spreading simulated events (lower right). **f and g**, Determination of average diffusion constants based on the widening of superimposed single events. **f**, Average of all 273 events with multiple Syp1-2xpHl for different time points. Scale bars 200 nm (x, y) and 0.2 a.u. (z). **g**, Widths (variance) of Gaussian fits to centered and averaged events in **f** for the first 10 frames (100 ms, solid blue line). A linear fit (dashed blue line) yields a diffusion coefficient of 0.1208 µm^2^/s for events with multiple Syp1-2xpHl. Monte Carlo simulations of four pHl with the apparent diffusion constant as input were performed and serve as a control. Widths of Gaussian fits to the average of 1000 simulated events (grey solid) can be fit with a line (grey dashed) yielding a diffusion constant of 0.1209 µm^2^/s. Fitting with a Gaussian function can only be performed on averaged signals, as fast spread of few particles may lead to asymmetrical intensity distributions. **h**, Histograms of spread values of 273 events from radial kymograph analysis (solid) and of 1000 simulated events (dashed) using the diffusion constant determined in **g**. The median spread values are 0.1407 µm^2^/s for measured and 0.1626 µm^2^/s for simulated events. **i**, Histograms of apparent diffusion constants of patch diffusion (center of intensity) of all 273 measured (solid) and 1000 simulated events (dashed) with a median spread value of 0.0014 µm^2^/s for measured and 0.0036 µm^2^/s for simulated events (n = 11 regions, 3 biological replicates). Note that the histogram of simulated data was normalized to the integral of the histogram of measured data.

### Optical single SV recordings reveal rapid dispersion of SV proteins post fusion

Do SV proteins remain in part clustered after exocytosis, thereby facilitating formation of endocytic structures or are they re-clustered at sites of endocytosis? To what extent might self-assembly forces between SV proteins figure in maintaining the RRetP? To this end, we examined the fate of several SV proteins post fusion using live-cell TIRFM imaging of xenapses expressing pHl fusion variants of a number of SV proteins, differing in size, the number of transmembrane domains as well as the abundance in SVs. As indicated above, we restricted analysis at first to exocytosis events with photon counts corresponding to at least two pHl-tagged proteins per SV. In particular, we investigated the postfusion fate of the tetraspan and second most abundant protein Syp1 (about 30 copies per SV ^48^), which binds the SV SNARE Syb2 and is important for clearance of the AZ ^49^, as well as of pHl-tagged versions of the calcium sensor Syt1 (10 – 15 copies per SV ^48^), the V0c subunit ^50^ of the large vacuolar ATPase (1 – 2 copies per SV ^48^) and of the vesicular glutamate transporter VGlut1 (10 – 15 copies ^48^). The high amount of surface-stranded molecules for Syb2-pHl overexpression ^9^ precluded such analysis for Syb2. A time-lapse series of a single exocytosis event with 7 Syp1-2xpHl clearly shows rapid radial dispersion of the molecules within 400 ms (Fig. 5a). However, in other examples with at least 4 pHl molecules dispersion is apparently absent (Fig. 5b, 2^nd^ event: columns 3 and 4). Gaussian fits cannot be applied for quantification of dispersion because of the low quantity of molecules moving. Thus, in order to quantify dispersion on this ‘mesoscale’ with only 4 molecules moving, we decided to plot radial kymographs around the origins of AP-triggered release events using slightly smoothed movies (Fig. 5b, 2^nd^ and 4^th^ column; wavelet filtered, cf. Methods). Kymographs are a vivid visualization of intensity changes over time and space. An event that spreads over time will be localized within a small annular distance first, while its intensity stretches towards larger annular distances for later times (Fig. 5d, top, suppl. Fig. 5.1a,b). Kymographs could be fit at each time point by a simple logistic function 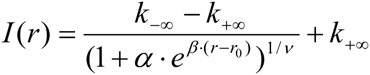, where *r* is the annular distance and *r*_0_ a horizontal shift, *k*_−∞_ and *k*_+∞_ denote the maximum and minimum intensities, *β* the growth rate, *α* a scaling factor, and *ν* determines the shape of the logistic function. The radius of half decay for each time point was determined from the logistic fit (suppl. Fig. 5.1c) and plotted as squared value against time (Fig. 5d, bottom). We restricted our analysis to the first 100 ms upon release, because at later times partial bleaching of released molecules might confound the dispersion analysis. Two extreme examples of dispersion behavior are shown in Fig. 5b: an event displaying rapid radial dispersion of pHl molecules and an event with no visible dispersion. Do events with no discernible dispersion correspond to ‘kiss-and-run’ events, or do they merely reflect cases, where the few released molecules during their random walk happen to stay close to the release site for the first 100 ms? To test this hypothesis, we performed Monte-Carlo simulations of a purely diffusional 2D random walk (cf. Methods). The only free parameter in these simulations is the diffusion constant, which we determined from our data. To this end, we centered all recorded fusion events in space and time (Fig. 5f). This compound movie now could be well fit at every time point by a 2D Gaussian function, and a plot of the fit variance against time could be fit with a straight line with a slope of D = 0.1208 µm^2^/s, indicating free diffusion for the first 100 ms (Fig. 5g). With this diffusion constant we performed 1,000 random walks of four particles deposited in the center at time zero and added to these images the measured camera photon shot noise and Gaussian noise, as experimentally determined for our measurement setup (cf. Methods). Examples of such simulated random walks are not distinguishable from experimental data (Fig. 5c, columns 1 and 3, or smoothed, 2 and 4, respectively). When subjecting both, experimental and simulated images to the same radial kymograph and logistic fit analysis, we could derive for every measured and simulated fusion event an individual spread parameter in μm^2^/s from a linear regression to the respective plot of the squared radii of half decay versus time. Histograms of these dispersion coefficients yield nearly identical skewed distributions for experimental and simulated fusion events (Fig.5h). Thus, most, if not all fusion events showed rapid dispersion by free diffusion. Even those events, which do not show overt dispersion within 100 ms, i.e. those events within the first bin of the histogram can be explained by free diffusion. Nevertheless, could some of these events represent ‘kiss and run’ fusion? To this end, we repeated above experiments and analysis in xenapses expressing the SV proteins Syt1-pHl, V-ATPase subunit V0c-pHl, or VGlut1-pHl (suppl. Fig. 5.2). Although SVs harbor only 1 – 2 copies of the V-ATPase per SV we were able to detect fusion events with more than 4 pHl molecules.

In summary, all pHl constructs tested showed free diffusional dispersion with diffusion constants close to 0.1 µm^2^/s, in the range expected for transmembrane proteins ^51^, but about four times lower than free diffusion of syb2-pHl in axonal membranes (suppl. Fig. 5.3). Given that the diffusion constants differ for each type of protein underscores the notion that SVs do not undergo ‘kiss-and-run’, but instead fully collapse after fusion. In addition, this finding rules out any patch diffusion, where SV cargo proteins remain clustered after fusion, as has been suggested in the first biological study capitalizing on the then new STED nanoscopy ^15^. We assessed patch diffusion in our data by determining the position of the center of mass or intensity for every time point and plotting the mean square displacement of those positions versus time for every event with at least two Syp1-2xpHl molecules. Fit of a regression line yields the diffusion constant of patch diffusion during the first 100 ms for every fusion event (Fig. 5i). The histogram of diffusion constants shows a very narrow distribution with an average value of 0.0014 µm^2^/s. With *x*^2^ = 4*Dt* for 2D diffusion this would correspond to a diffusion distance of 24 nm, i.e. just half a SV diameter, which would result in massive STD because of clogging ^41^.

Our finding of rapid diffusional dispersion, however, also implies that outside the AZ cargo cannot be re-clustered and re-assembled into retrieval-ready patches simply by self-assembly forces, as interactions among SV proteins (e.g. Syp1/Syb2 co-oligomers) are rather weak (reviewed in ^52^). Our findings thus favor a scenario, where released SV proteins are recaptured by specific adaptor proteins and these adaptor-cargo complexes are cross-linked by Clathrin triskelia. Therefore we examined whether patches of the RRetP are decorated with Clathrin.

### The RRetP counterbalances the RRP and recaptures SV proteins outside the AZ

As xenapses are strongly adhering to the polymer nano-brush by transsynaptic cell adhesion complexes, we attempted to isolate the cell membrane cortices from xenapses for ultrastructural analysis by ‘unroofing’ using short pulses of sonication. Membrane sheets were prepared at 4° C to prevent endocytosis and minimize exocytosis and were chemically fixed and coated with 2 nm platinum/carbon. Subsequent SEM of platinum replicas of the inner membrane surfaces revealed numerous pre-assembled Clathrin-coated structures (CCS) (Fig. 6a). Quantification of the density of these Clathrin patches yields an average of 8.8±3.7 CCS per µm^2^ (17 xenapses), i.e. about 80 CCS per xenapse (with typical footprint area of 7 to 10 µm^2^) and about twice the number of SVs found in the RRP (Fig. 4h,i). Electron micrographs of vertical thin sections of unroofed xenapses confirm that these CCS are formed on bouton membranes at the peri-active zone (Fig. 6b). Freeze-fracture replicas of the inner leaflet of the xenapse plasma membrane facing the coverslip reveal dimples matching in size the domed CCS seen in unroofed xenapses that are enriched with 4 to 8 particles (Fig. 6c), as also seen in frog neuromuscular junctions ^14^. In small synapses we found previously, that a train of 70 to 80 APs is needed to deplete the RRetP ^9^, compared to 40 to 50 APs needed for RRP depletion, showing, that the ratio of RRetP and RRP sizes is similar both in small hippocampal boutons and xenapses.

**Figure 6.**
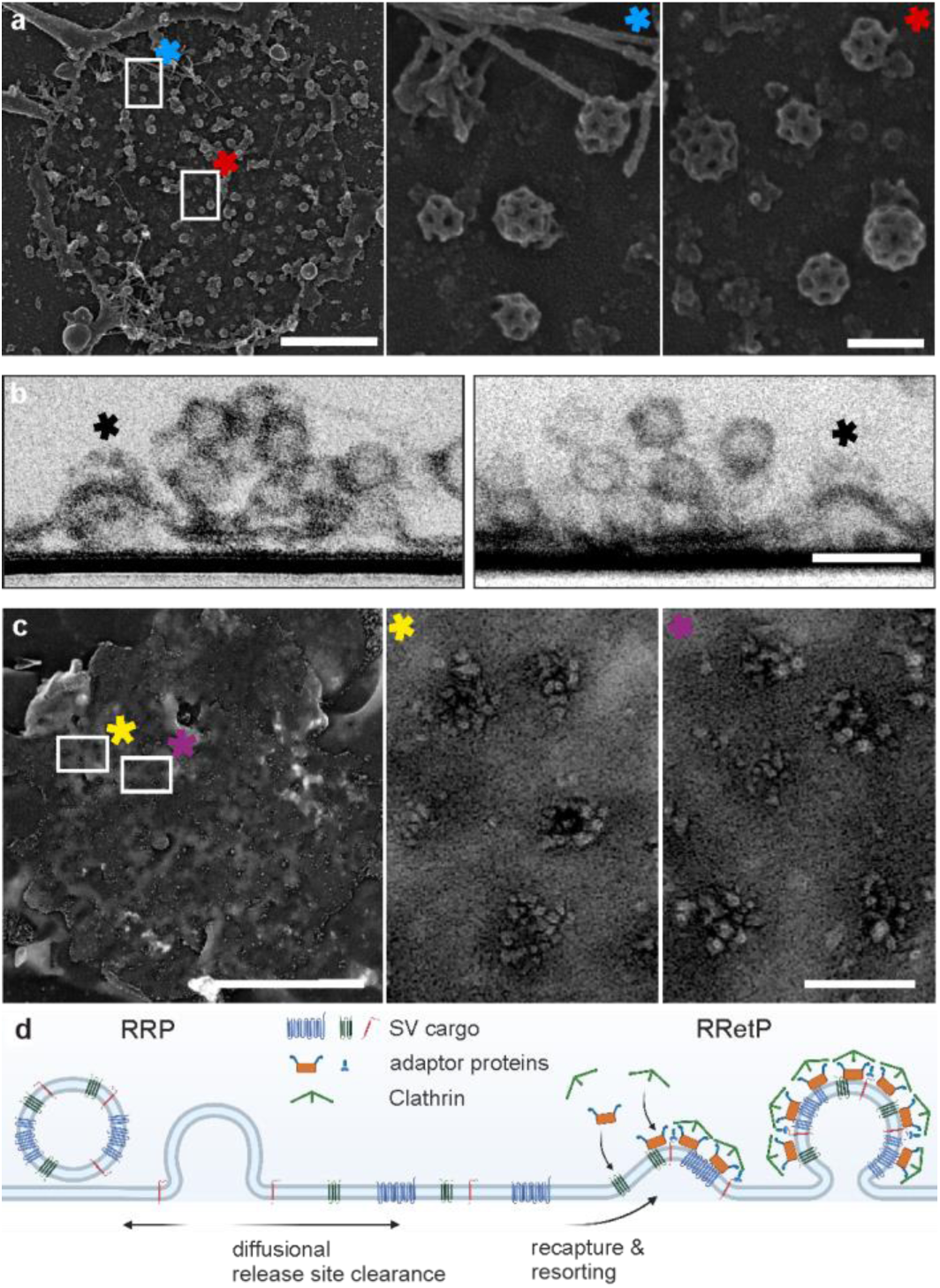
The RRetP counterbalances the RRP in size and recaptures SV proteins outside the AZ. **a**, Overview SEM image (left) of Pt replicas of unroofed xenapses showing numerous CCSs (8.84 ± 3.6 per µm^2^), remaining clusters of SVs, and actin filaments. Scale bar 1 µm. Asterisks and rectangles in the overview image mark the locations of the detailed views (center and right). Scale bar 0.1 µm. **b**, Electron micrographs of vertical thin sections of unroofed xenapses confirming that Clathrin structures (marked with asterisk) are formed on bouton membranes at the peri-active zone. **c**, Overview of TEM image (left) of Pt replicas from the inner leaflet of the bottom membrane of LN2-frozen and freeze-fractured xenapses showing CCS sized dimples. Scale bar 1 µm. Asterisks and rectangles in the overview image mark the locations of the detailed views (center and right). 4 to 8 large integral membrane proteins are visible indicating sorting of exocytosed and dispersed SV cargo. Scale bar 0.1 µm. **d**, Schematic summarizing the results. Rapid diffusion of vesicle proteins after fusion clears the release site. Cleared proteins are recaptured and resorted by adaptor proteins and Clathrin into patches of the RRetP outside the AZ, which explains the non-identity of molecules released and retrieved.

## Discussion

A number of biomimetic tools to study synapse formation have been developed that induce synapses in a more defined manner compared to regular dissociated neuronal cultures ^26–28^. Here we introduce a novel pre-synaptic culture, the xenapse, which precisely defines the formation of the synaptic structure in two dimensions directly on the substrate. Thorough structural and physiological characterization revealed, that xenapses share many features with small synapses, including active zone, RRP and RRetP sizes, phasic responses to trains of APs and synchronous SV exocytosis followed by fully compensatory endocytosis.

A powerful evidence in favor of SV full-collapse fusion over ‘kiss-and-run’ is the EM data pointing to Clathrin-mediated endocytosis (CME) being the major mechanism for SV protein recapture and recycling ^12,53^. As CME might be too slow for fast SV recycling, we and others have suggested that a pre-sorted and pre-assembled pool of SV proteins, the RRetP, on the presynaptic membrane might support a first wave of fast CME ^6,9,14,18^. The loss of vesicular molecular identity resulting from SV flattening necessitates the repackaging of the cargo, achieved by specific adaptor proteins that link Clathrin coat formation to the concomitant selection and concentration of cargo proteins. Although all experiments in this study were performed at room temperature, we observed endocytosing vesicles to be drawn from the RRetP by CME also at physiological temperature in response to single APs, i.e. UFE conditions (data not presented here).

Interestingly, interference with the clearance of SV proteins after fusion away from the active zone and their retrieval at the periactive zone results in rapid STD, likely induced by clogging of release sites ^41,54^. This finding necessitates a mechanism for fast clearance of SV proteins post fusion within a few 10 ms, as SVs can be reprimed after depletion within 50 – 100 ms ^45^. Diffusion constants of about 0.1 µm^2^/s which we measured here for SV protein clearance fulfill this requirement. With *x*^2^ = 4*Dt* for 2D diffusion, SV proteins would travel on average a distance of 200 nm within 100 ms. This is consistent with the analogous analysis of evanescent wave microscopy images of secreting chromaffin cells which revealed fast dispersion of Syb2-EGFP with a diffusion constant of 0.2 µm^2^/s during dense-core granule exocytosis, except for a few events representing transient fusion without full collapse ^55^.

However, a FRAP (fluorescence recovery after photobleaching) analysis of free diffusion of Syb2-pHl in axonal membranes, which unlike the other pHl constructs displays a sufficiently large surface pool, yielded an average diffusion constant of 0.38 µm^2^/s (suppl. Fig. 5.3), i.e. fourfold higher than in the AZ. This discrepancy between free diffusion of surface-stranded SV proteins in axons and the AZ adjacent regions in boutons could arise from two effects: first, AZs in EM micrographs appear as electron-dense thick endings next to the PM, implying a dense network of large scaffold proteins ^56^. This dense matrix on the inner surface of the PM should slow diffusion of transmembrane proteins within the PM due to increased tortuosity. Second, recapturing of cargo outside the AZ by Clathrin-coated RRetP patches will decrease the apparent diffusion constant.

For events with at least two Syp1-2xpHl molecules, evoked by a single AP, we could not detect clear dispersion in only about 10 % of these events (Fig. 5h). The abundance of not-dispersing events, however, was comparable to that predicted by Monte-Carlo simulations of free diffusion of four diffusing molecules during the first 100 ms (Fig. 5h). In principle, some of these events might represent true kiss-and-run which is in line with previous cell-attached capacitance measurements of single exo- and endocytic events at pituitary nerve terminals ^57^ and calyces of Held ^58^. These revealed only rarely transient fusion with measurable fusion pore conductance which could reflect kiss-and-run. Alternatively, such apparent non-dispersing of cargo could represent events where proteins remained clustered post fusion. In earlier high-resolution STED microscopy clusters of Syt1 were detected by immunofluorescence on the PM after stimulation ^15^. These were taken as evidence that the SV protein syt1 did not disperse during recycling but remained clustered although no live-cell imaging was performed as proof. Also, as has been noted elsewhere, the use of secondary immunodetection itself could have led to Syt1 clustering in these STED images ^18^. However, this earlier observation of surface Syt1 patches is fully compatible with the notion of fast recapturing of dispersed SV proteins into an RRetP. Recently, cryo-electron microscopy of native SVs identified Syp1 as an interaction partner of the V-ATPase in a 1:1 stoichiometry ^59,60^, which supports the notion of a role of self-assembly in SV biogenesis. However, given the large difference in copy numbers per SV for these two proteins, this interaction, even if with high affinity, cannot account for SV proteins remaining clustered in an 80-nm patch after fusion. Our measured diffusion constant of 0.0014 µm^2^/s for patch diffusion implies that a lack of dispersion would cause clogging of release sites. We could indeed previously show that accumulation of syt1 in AZs, as seen in STED images after acute endocytosis block immediately results in strong STD ^41^. Furthermore, EM images show a very compact organization of SVs at the membrane ^61^, consistent with functional data showing vesicles to be rather immobile even during synaptic activity ^62,63^. Likewise vesicles appear to undock only once per about two minutes at hippocampal synapses ^64^. Encircled by other vesicles a vesicle transiently fusing may not be easily pushed aside or undocked thereafter but would have to be re-acidified and primed for reuse in place, which takes however up to 5 s ^65^. Hence, with ‘kiss-and-run’ as major mechanism the refractory period of a given release site would be about 5 s. This is a much longer timescale than that for translocation, docking, and priming at hippocampal synapses (fast component of 1.5 s) ^66^ or also in the Calyx of Held ^40^. Thus, our results bolster existing data to establish that most if not all SV fusion events result in full collapse and rapid dispersion of vesicle proteins in the plasma membrane (Fig. 6d). This mechanism ensures that vesicle proteins are rapidly and efficiently cleared from the release site, such that its intricate molecular organization and integrity is not distorted.

The highly efficient confinement of SV proteins within the bouton membrane by the RRetP resolves the apparent contradiction between free redistribution of pHl-tagged SV proteins within axonal membranes and their enrichment in the bouton membrane, typically observed also in small synapses ^49^. Additionally, an effective barrier for lateral diffusion in the membrane may confine vesicle proteins to the bouton. Replicas of unroofed xenapses indeed show a rather dense network of cortical actin filaments in-between areas of Clathrin-coated structures (Fig. 6a). The swift and efficient re-capturing and re-clustering of rapidly dispersing SV constituents by the extended RRetP provides a diffusional sink, efficiently reducing back-diffusion of exocytosed SV proteins into the AZ, and thus prevents release-site clogging and resulting STD ^49^. Such mechanism of fast SV collapse, rapid diffusional dispersion, and efficient recapturing outside the AZ, as reported here, allows CNS synapses to transmit frequency-coded information with broad bandwidth.

## Materials and Methods

Extended methods at end of this file.

Supplementary Information is available for this paper.

## Acknowledgements

We thank K. Tkotz and A. Roetrige for excellent technical support. This work was supported by grants of the Deutsche Forschungsgemeinschaft (SFB 944 P5 and SFB 1348 A02 to J.K. and SFB 1348 A03 to M.M.).

## Author information

The authors declare no competing financial interests. Correspondence and requests for materials should be addressed to J.K.

## Author contributions

J.L. S.K, M.K. and J.K. conceptualized the study and designed the experiments; J.L., J.D., L.P.S., G.N. J.H. and P.S. conducted and analyzed the imaging experiments; N.G, Y.T, and U.K. conducted and analyzed EM experiments, C.R. and M.M. designed and cloned Halo-Nlgn1 extracellular domain, J.L. and J.K. wrote software for data analysis; and J.K., S.K., and M.K. supervised work and wrote the paper with contributions from all authors.

## Data availability

The data that support the findings of this study are available from the corresponding author upon reasonable request.

## Code availability

Matlab Code is available on GitHub (https://doi.org/10.5281/zenodo.13849665).

## Methods

### Animals

Breeding of C57BL/6N mice was conducted at the Central Animal Experimental Facility of the University Hospital Muenster according to the German Animal Welfare guidelines (permit number AZ81–02.05.50.20.019, issued by ‘Landesamt für Natur, Umwelt und Verbraucherschutz NRW’, Duesseldorf, Germany). Both male and female mice were used for primary neuronal cultures.

### Plasmids and molecular biology

pCAAGS-vGlut1-pHluorin (VGlut-pHl) was a gift from R.H. Edwards ^1^. pcDNA3-Syp1-pHluorin 2x (Syp1-2xpHl) was a gift from Stephen Heinemann and Yongling Zhu (Addgene plasmid # 37004; http://n2t.net/addgene:37004; RRID:Addgene_37004). The plasmids encoding Synaptotagmin1-pHluorin (Syt1-pHl) and V0c-pHluorin (V0c-pHl) have been described elsewhere ^2,3^. To visualize endogenous Munc13, the HITI (homology-independent targeted integration) based ORANGE (open resource for the application of neuronal genome editing) approach was used to tag endogenous Munc13 with EGFP ^4^. This corresponding vector contains the entire cassette for knock-in and was a gift from Harold MacGillavry (Addgene plasmid # 131498; http://n2t.net/addgene:131498; RRID:Addgene_131498).

Neuroligin-1 extracellular domain (Nlgn1ECD) fused to human Fc-tag has been interspersed with Halo protein followed by an HRV 3C protease cleavage site to obtain purified Nlgn1ECD-halo protein. Mutagenesis was used to generate the specific Nlgn splice variant and to insert HRV 3C recognition site, with the mutagenesis being performed with QuikChange method (Stratagene/Agilent Technologies) using complementary primer pairs (forward primer and silent restriction site for recognition are in parentheses). The resulting vectors were sequenced (GATC Biotech AG, Konstanz, Germany). pCMV-Nlgn1(+A-B)-Fc was generated as splice insert B negative variant of pCMVIG-NL1(+A+B) ^5^ using forward primer (5’-GCT GAC TTT ATC CCA TTA TTC TGA AGG ACT TTT TCA ACG AGC AAT AGC -3’, -BstEII). A 3C protease cutting site was inserted by Quikchange using forward primer (5’-GAT AAG CTT GCA TGC CTC GAG GTA CTA TTC CAG GGA CCG CTG CAG GTC GAC TCT AG-3’) to obtain pCMV-Nlgn1(+A-B)-Cys-3Cpro-Fc. Halo-tag from IfNAR-Halo ^6^ was amplified by standard PCR with primers (5’-CT ATC GAT AAG CTT GCA GAA ATC GGT ACT GGC TTT CCA TT-3’, +ClaI, +HindIII; 5’-C CTC GAG ACC GGA AAT CTC CAG AGT AGA CA-3’, +XhoI) and ligated into pCMV-Nlgn1(+A-B)-Cys-3Cpro-Fc at BamH1 and XhoI sites flanking the cys to yield pCMV-Nlgn1(+A-B)-Halo-3Cpro-Fc (’Nlgn1-Halo’). For protein expression in mammalian cells, a NotI/XbaI-digested fragment containing Nlgn1(+A-B)-Halo-Fc was inserted into pTT/pPLE(+) vector. NotI and XbaI sites were introduced using forward primers (5’-GAATTCAGATCTGCGGCG GCC GCCACCATGGCACTTCCC-3’ and 5’-GTGCGACGGCCG CTC TAG AGACTCTAGGGGTG-3’).

### Protein expression and purification

pTT/pPLE(+)-Nlgn1(+A-B)-Halo-Fc was transiently transfected into HEK-INVcells in 1 L serum-free medium (done by InVivo Biotech Services). Fc-Fusion proteins that were secreted into FCS-free medium were bound to rmp Protein A Sepharose^®^ Fast Flow beads (50 µL slurry beads per 100 mL medium, Sigma), precipitated (30 s, 11.000 g), washed with HEPES buffer (10 mM HEPES, 150 mM NaCl, pH 7.5) and digested (4 °C, overnight) by 6 units of Pierce™ HRV 3C Protease (Thermo Fisher Scientific) in a volume of 500 µL. After removal of the his-tagged protease with Ni-NTA agarose beads (Qiagen) in HEPES buffer (10 mM HEPES, 300 mM NaCl, pH 7.5), the supernatant was syringe filtered using Whatman™ 0.2 µm Cellulose acetate filters (GE Healthcare Life Sciences) and concentrated via Pierce™ 3K PES Protein Concentrators (Thermo Fisher Scientific) to approximately 1.0 mg/mL. The final product is the soluble fusion protein Nlgn1 ECD-Halo. (residues 46-663, *rat*, NLGN1_RAT isoform 3, Q62765-3) with splice insert A but not B (residues 299-NRWSNSTKG-307) and a C-terminal Halo7-tag (Acc no. AY773970).

Sequences coding for superecliptic pHluorin and EGFP were cloned into pET-24a(+) vectors (Novagen) containing a C-terminal hexahistidine affinity tag sequence and subsequently transformed into BL21(DE3)pLysS competent cells (Novagen). Target proteins were purified based on immobilized metal affinity chromatography using a Ni^2+^-loaded HiTrap chelating column (GE Healthcare) connected to an ÄKTA prime plus chromatography system (GE Healthcare). Elution was performed with a linear imidazole gradient starting from 0 mM imidazole in buffer A (20 mM HEPES, 150 mM NaCl, pH 8) to 500 mM imidazole in buffer B (20 mM HEPES, 150 mM NaCl, 500 mM imidazole, pH 8). Eluted fractions were pooled based on the UV absorbance (measured at 280 nm) in the chromatogram, concentrated using Amicon Ultra-4 centrifugal filters (UFC801024, Millipore) and further purified by size exclusion chromatography using a HiLoad 16/600 Superdex 75 pg column (GE Healthcare) equilibrated with buffer A at a typical flow rate of 0.5 ml/min.

### Preparation of functionalized PLL-PEG derivatives

Poly-L-lysine (PLL) grafted with polyethylene glycol (PEG)-HaloTag ligand (HTL) (PLL-PEG-HTL) for specific immobilization of Halo-fusion proteins on surfaces was prepared as published ^7^. PLL grafted with PEG-Methoxy (PLL-PEG-OMe) was purchased from SuSoS AG.

### Patterning of glass coverslips via microcontact printing (µCP)

Structured Silicon wafers for the casting of µCP stamps were custom made (Nanotechnology and Devices/NT&D). Polydimethylsiloxane (PDMS) stamps were prepared from Sylgard 184 silicone elastomer (Dow Corning) as described elsewhere ^8,9^. 18 mm microscope cover glasses (Glaswarenfabrik Karl Hecht GmbH) served as substrates and were cleaned with oxygen plasma in a FEMTO plasma cleaner (Diener Electronic). Inking of the stamps was performed for 45 minutes with PLL-PEG-HTL followed by washing of the stamps with deionized water. Dry stamps were brought into contact with cleaned coverslips for 1 minute to transfer the ink to the substrate. After peeling off the stamps, the micropattern was backfilled for 1 minute with PLL-PEG-MeO which, with its net negative charge, repels cell growth outside the stamped pattern. After washing the coverslips with deionized water, Nlgn1-Halo was applied for 1 hour in a humidified CO_2_ incubator. After removing excess Nlgn1-Halo by applying culture medium supplemented with 10% FCS, coverslips were placed in a 12 well plate with a glial feeding layer and preconditioned neuronal growth medium in a CO_2_ incubator. Preparation of primary neurons was performed a few hours after the latter step.

### Neuronal cell culture and transfection

Primary murine hippocampal neurons were prepared from brain of newborn pups (P0). The hippocampi were dissected in ice-cold artificial cerebrospinal fluid (ACSF, 137 mM NaCl, 26 mM NaHCO_3,_ 8 mM Na_2_HPO_4,_ 2.7 mM KCl, 1 mM KH_2_PO_4,_ 1 mM MgCl_2_, 30 mM D-glucose), and digested with trypsin (1 mg/mL trypsin (Sigma-Aldrich) and 0.25 mg/mL DNase (Sigma-Aldrich) in 137 mM NaCl, 5 mM KCl, 7 mM Na_2_HPO_4_, 25 mM HEPES, pH 7.4) for 20 minutes at 37 °C. Trypsin was then inhibited by twice being washed with plating medium ((MEM, Gibco) supplemented with 5 g/l glucose, 200 mg/l NaHCO_3_, 100 mg/l transferrin (Calbiochem), 25 mg/l insulin (Sigma), 10% fetal bovine serum (Sigma-Aldrich), and 2 mM L-glutamine (Sigma-Aldrich)). The digested tissue pieces were triturated with fire-polished siliconized Pasteur pipettes in plating medium containing DNase (0.25 mg/ml). The cell suspension was diluted with growth medium (NBA (Gibco) supplemented with 2% B27 (Gibco), 2 mM Glutamax (Gibco) and 0.1% penicillin/streptomycin (Gibco)). Finally, neurons were plated onto 18 mm micro-patterned glass coverslips (35,000 cells per coverslip). Coverslips were placed on silicon feet on top of the glial feeding layer in 12 well plates. To inhibit glial cell proliferation, 2 µM cytosine arabinoside (AraC, Sigma-Aldrich) was added to the culture medium at DIV 3-4. Axonal varicosities, the xenapses, could be seen on the patterned substrate after about 24 hours in culture. Dense continental hippocampal cultures and astrocytes for feeding layers were prepared as described previously ^10^. Transfection was done at DIV 5 following a modified calcium phosphate transfection protocol ^11^. Neurons were imaged on DIV 8-21.

### Surface analysis of PDMS stamps and micropatterned coverglasses with AFM

The topography of microstructured patterns was measured with a NanoWizard II AFM (JPK Instruments) using the intermittent contact (IC) mode in air at room temperature (RT). To this end, the Point Probe Plus (PPP)-NCH cantilever tips (Nanosensors) with spring constants between 10 N/m and 130 N/m were oscillated slightly below the resonance frequency of 270 to 300 kHz to allow for height imaging, at a scan rate below 0.3 Hz, while the axial scan range was set to 5.85 µm. Image analysis and processing were carried out using the JPK image processing software as well as GIMP (https://www.gimp.org).

### Immunostaining of micropatterned coverglasses

After µCP, the substrates were rinsed in PBS, incubated in blocking buffer (3% bovine serum albumin (BSA) and 0.1% cold-water fish skin gelatine (FSG, Sigma-Aldrich) in phosphate buffered saline (PBS, Sigma-Aldrich)) for 1 hour, followed by incubation with a primary antibody against the extracellular domain of Nlgn1 (rabbit polyclonal, dilution 1:200, Synaptic Systems). After washing three times with PBS, substrates were incubated with a secondary antibody (goat-α-rabbit Alexa488, dilution 1:1000, Thermo Fisher Scientific) for 1 hour and finally washed again three times with PBS.

### Immunostaining of xenapses

Xenapses were washed twice with prewarmed isoosmotic Hank’s buffered salt solution (HBSS, Gibco) and fixed with 4% (v/v) paraformaldehyde (PFA, Sigma-Aldrich) in PBS containing 4% (w/v) sucrose (Sigma-Aldrich) for 20 minutes at RT. After washing with PBS, cells were incubated with 0.1 M glycine (Carl Roth) for 20 minutes to quench remaining PFA. Then, cells were blocked and permeabilized in blocking buffer (3% (w/v) BSA (Sigma-Aldrich), 0.1% (w/v) FSG, and 0.2% (w/v) saponin in PBS) for 1 hour at RT. Cells were incubated with primary antibodies in blocking buffer overnight at 4 °C. Then, cells were washed with washing buffer (0.2% (w/v) BSA and 0.05% (w/v) saponin in PBS) five times and incubated with secondary antibodies in blocking buffer for 4 hours at RT. Finally, cells were washed five times with washing buffer, once with PBS and mounted in Mowiol. The following primary antibodies were used: α-tau-1, guinea pig polyclonal (1:200 dilution, Synaptic Systems), α-MAP-2 guinea pig polyclonal (1:100 dilution, Synaptic systems), α-Synaptophysin1, rabbit polyclonal (1:250 dilution, Synaptic Systems). For detection, the following secondary antibodies were used: goat-α-rabbit AlexaFluor488 (1:1000 dilution, Thermo Fisher Scientific) and goat-α-mouse AlexaFluor594 (1:1000, Thermo Fisher Scientific).

### Sample preparation for TIRF-dSTORM

Xenapses (DIV 10-18) were washed twice with pre-warmed iso-osmotic HBSS and fixed with 4% (v/v) PFA (Sigma) in PBS containing 4% (w/v) sucrose (Sigma) for 20 minutes at RT. Subsequently, they were washed with PBS and free aldehydes were quenched in 0.1 M glycine (Carl Roth) in PBS for 10 minutes. Next, samples were incubated in blocking solution (3% (w/v) BSA (Sigma-Aldrich) and 0.1% saponin (Sigma-Aldrich) in PBS) for 1 hour at RT, followed by incubation with primary antibodies in blocking solution overnight at 4°C with gentle rocking. Next, they were washed five times for 5 minutes each in washing buffer (0.2% (w/v) BSA and 0.01% (w/v) saponin in PBS). Subsequently, they were stained with Alexa-labelled secondary antibodies for 1 hour followed by washing them five times for 5 minutes each with the washing buffer. A primary antibody against Bassoon (mouse monoclonal, 1:500 dilution, Enzo) and a secondary antibody conjugated to Alexa 568 (goat-α-mouse Alexa Fluor 568, 1:1000 dilution, Thermo Fisher Scientific) were used. Next, the samples were post-fixed in 4% PFA with 0.2% glutaraldehyde (Sigma-Aldrich) for 20 minutes. After post-fixation, samples were sequentially washed with PBS for 5 minutes, quenched in 0.1 M glycine (in PBS) for 10 minutes, and again with PBS for 5 minutes. To correct for x-y drift during imaging, fiducial markers (fluorescent 0.1 µm TetraSpeck microspheres, Invitrogen) were added to the sample in a 1:2000 dilution in PBS, and incubated for 20 minutes at RT followed by two washes with PBS. Samples were stored in PBS.

### TIRF-dSTORM imaging

Imaging was performed on an inverted Nikon Eclipse Ti microscope, equipped with a TIRF illumination condenser, a 100x / 1.49 NA Apo TIRF oil immersion objective (Nikon) and a 561 nm TIRF filter set (ZET 561/10, zt 561/650/50 ET, AHF). A sCMOS camera (Prime 95B, 11 x 11 µm^2^ pixel area, Photometrics) was side mounted on the setup and a laser light engine (Sole-6, Omicron-Laserage Laserprodukte GmbH) with high-speed analog and digital modulation was used to control a 561 nm diode-pumped solid-state laser (Jive, 150 mW output, Cobolt AB/Hübner Photonics GmbH). Image acquisition was controlled by µManager software 1.4.19 ^12^. Measurements were performed in photo-switching solution which was based on a buffer containing 200 mM piperazine-N,N’-bis(2-ethanesulphonic acid) (PIPES, Carl Roth) at pH 7.2 and 40% (w/v) glucose (Carl Roth), freshly supplemented with 100 mM *β*-mercaptoethylamine (also known as cysteamine, Sigma-Aldrich), 143 mM *β*-mercaptoethanol (Sigma), 0.8 mg/ml glucose oxidase (Sigma-Aldrich) and 0.08 mg/ml catalase (Sigma-Aldrich). Fluorescent molecules were converted into a long-lived dark state with a 405 nm diode laser (PhoxX, 60 mW output, Omicron-Laserage Laserprodukte GmbH) at an irradiation intensity of 9.6 ± 0.3 W/cm^2^ on the sample. Once optimal blinking was achieved, the illumination mode was swapped back to TIRF mode and a 2×2 binned image time lapse (pixel size = 220 nm) was acquired with an exposure time of 50 ms, an acquisition frequency of 20 Hz and a total of 20,000 images.

### dSTORM reconstruction and cluster analysis

Reconstruction was performed using the ImageJ plug-in ThunderSTORM ^13^. Single molecule coordinates were determined by fitting a 2D Gaussian function to the single molecule intensity profiles using the Levenberg-Marquardt algorithm. Lateral drift was corrected by tracking fiducial markers using a trajectory-smoothing factor of 0.05. Localizations associated with a standard deviation of the Gaussian fit higher than 200 nm were filtered out before image reconstructions. The molecule coordinates were exported to SR Tesseler ^14^, where Voronoi tessellation was performed on all detections. Clusters (called objects in SR Tesseler) were segmented using an average localization density factor of 2 (over a spatially uniform localization distribution in the same area), a cut distance of 30 nm, a minimum cluster area of 0.008 μm^2^ and a minimum of 100 localizations per cluster. For each cluster identified SR Tesseler returned cluster dimensions such as area, number of detections and diameter.

### STED microscopy of endogenous Munc13-1 in xenapses

Cells were transfected on DIV 4 with pORANGE Munc13-1 GFP KI using the Effectene transfection reagent (Qiagen). On DIV 18, xenapses were fixed and immunostained for Bassoon using the same procedure as for dSTORM sample preparation without the post fixation step. To amplify the EGFP signal of endogenous Munc13, an Abberior Star 635 P-labelled nanobody targeting EGFP (1:200, Abnova) was added while incubating with the secondary antibody. Bassoon was labelled using a primary antibody targeting its N-terminal domain (mouse monoclonal, 1:500 dilution, Enzo) and an Alexa 594-labelled secondary antibody (goat-α-mouse Alexa Fluor 594, Thermo Fisher Scientific, 1:1000 dilution). The coverslips were mounted on glass slides with Mowiol (Sigma-Aldrich) supplemented with 250 mg/ml 1,4-diazabicyclo 2.2.2 octane (DABCO, Sigma-Aldrich) and STED microscopy was performed after adequate curing for two days at 4 °C. Imaging was performed using a Nikon Ti-E microscope with an attached STEDYCON scanner (Abberior Instruments) controlled by STEDYCON Smart Control software. A 100x / 1.45 NA oil immersion objective (Nikon) was used for imaging. Excitation lasers of 594 and 640 nm were used to sequentially excite Alexa Fluor 594 and Abberior Star 635 P dyes in confocal and STED mode. For the STED mode, a pulsed 775 nm depletion laser was used to create a xy depletion donut around the excitation spot. Signals were collected by avalanche photo diodes after passing a dual bandpass filter (537/35 nm and 617.5/19 nm), or a single bandpass filter (675/25 nm). A pixel size of 20 nm, a pixel dwell time of 10 µs, a time gating of 1 to 7 ns, a line accumulation of 2, and a depletion laser intensity of 90 - 100% were used for imaging.

### Live cell TIRF imaging

All experiments, unless otherwise stated, were carried out in standard buffer (140 mM NaCl, 2.5 mM KCl, 2.5 mM CaCl_2_, 1 mM MgCl_2_, 10 mM glucose, 10 mM HEPES, pH 7.4) at RT. NH_4_Cl solution was prepared by substituting 50 mM NaCl in standard buffer with NH_4_Cl. For Ca^2+^ imaging, xenapses were incubated with 5 µM Fluo5F-AM (Thermo Fisher Scientific) in osmolarity-adjusted physiological imaging buffer at pH 7.4 in a humidified CO_2_ incubator. After 10 minutes, the cells were washed three times. TIRF imaging of xenapses was performed on an inverted Eclipse Ti microscope (Nikon) equipped with a TIRF illumination unit, a 100x TIRF objective (NA=1.49, Nikon) and an Orca Flash 4 V2 sCMOS camera (Hamamatsu Photonics) with a pixel size of 65 x 65 nm^2^ at experimental conditions. Image acquisition was controlled by µManager software 1.4.19 ^12^. Epi-illumination was performed using a pE-100 white LED (CoolLED), TIRF illumination with a 488 nm diode laser (PhoxX, 200 mW output, Omicron-Laserage Laserprodukte GmbH). The optical filter set comprised a 2 mm thick ultra-flat dichroic mirror with cut-on wavelength of 491 nm (Chroma) and excitation as well as emission filters with bandpass wavelengths at 475/35 nm (Semrock) and 537/42 nm (AHF Analysentechnik AG). The penetration depth of the evanescent field generated was set to 80-90 nm. Neurons were stimulated by electric field stimulation (platinum electrodes, 10 mm spacing, 1 ms pulses of 50 mA and alternating polarity at 5-20 Hz) applied by a constant current stimulus isolator (World Precision Instruments). 10 µM 6-cyano-7-nitroquinoxaline-2,3-dione (CNQX, Tocris) and 50 µM D,L-2-amino-5-phosphonovaleric acid (AP5, Tocris) were added to prevent recurrent activity. Stimulation and imaging were synchronized using an Arduino Uno microcontroller unit (Arduino IDE 1.6.0, Arduino, www.arduino.cc). Frames were acquired with 100 Hz and an exposure time of 10 ms for single AP measurements.

### Analysis of live cell images

Live cell image analysis was performed using the 64-bit version of Matlab 2019a (MathWorks).

#### Peak detection

For peak detection, a template-based peak fitting algorithm was used, with a two-dimensional Gaussian function as approximation for the point spread function (PSF) as template. The Gaussian function was obtained by fitting the fluorescence signals of single fluorescent beads (TransFluoSpheres, 40 nm, ex./em. 488 /560 nm, Thermo Fisher Scientific). While shifting the position of the template, a fast Sum Square Difference (SSD) algorithm ^15^ accounts for the smallest deviation between template and PSF. Extension with a newer class of algorithms ^16^ enables subpixel resolution. Here, the template is resampled incrementally with higher subpixel resolution (0.1 pixel and 0.02 pixel, respectively), i.e. shifted by 0.1 pixel or 0.02 pixel in space.

The SSD is defined as

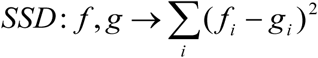

with the properties

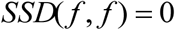

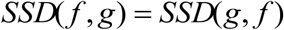

as a relation between function *f* and template *g*, which is zero in case of perfect match *g = f*. Because the SSD relation is symmetric, the template (Gaussian fit of the PSF) can be resampled instead of the image to achieve subpixel resolution. Thus, a new series of subpixel resolution templates are created with the template shifted by one subpixel in all directions. The algorithm was applied to a subregion of 9 x 9 pixel within both the normalized original image and the resampled templates as more than 90% of the theoretical PSF are included in this area. The subregions were excised around the brightest pixel of a segmented area found by *’à trous’* wavelet filtering ^17^. SSDs were calculated between this excised region of interest (ROI) and subpixel-shifted resampled versions of the PSF, and the minimum SSD was chosen for subpixel resolution localization. The minimization was done incrementally to reduce computer time, i.e. SSDs were calculated for templates shifted by 1 subpixel (= 0.1 pixel) against the ROI to yield an interim subpixel resolution localization, around which the templates with 0.02 pixel accuracy were shifted in a second step. The coordinates resulting from the second step were then stored as the molecular position. In the experiments for estimating RRP size all fusion events, i.e. also partially overlapping ones, had to be detected. To this end, we employed a deflation strategy as described ^18^ The template-based peak fitting algorithm is very robust even in the case of low signal-to-noise ratio or distorted PSFs. Although recorded fluorophores disperse after SV fusion, they could be fit reliably with the application of templates up to 1.5-fold broader than the system PSF.

#### pHluorin intensities

Intensities are given in number of photons (*N _photon_*) and can be converted from digital numbers (*DN*) according to,

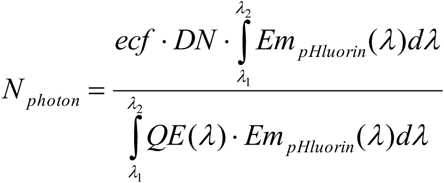

with the electron conversion factor *ecf*, the emission spectrum *Em _pHluorin_* of pHluorin within the respective detectable wavelength range λ_1_ - λ_2_ and the quantum efficiency *QE* of the camera. For determining the number of pHluorin molecules released per synaptic vesicle, purified pHluorin and for comparison EGFP were immobilized on coverslips by a polyacrylamide gel ^19^ at a concentration of 40 mM. Blinking events of the fluorophores were measured with maximum intensity of the 488 nm laser (9.6 ± 0.3 W/cm^2^) and an exposure time of 10 ms, followed by localization with the template based peak fitting algorithm, and subsequent integration of their intensities in a 9 x 9 pixel area.

#### Radial kymograph analysis

Annuli with increasing diameter were centered around the signal intensity maximum. Pixel intensities were then summed up and normalized for each ring to yield an averaged intensity per annular distance or per space unit. For each of the *n* time points, the intensity can be described as a function of the *m* radii by a logistic function ^20,21^ of the form

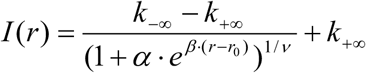

where *r* is the annular distance, *r_0_* a horizontal shift, *k_-_*_∞_ and *k_+_*_∞_ denote the maximum and minimum intensities, β describes the growth rate, α is a scaling factor for the exponential term, and ν determines the shape of the logistic function. While the overall intensity *I(r)* decreases over time as the particles diffuse out of the center, the area that the particles explore increases. The radius of this area can be described as that of full width at half maximum (FWHM). The apparent spread parameter *D_a_* was determined according to

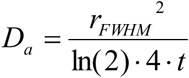

Notably, the distribution of spreading particles is not required to be symmetric for this analysis. This renders the radial kymograph a useful tool for analyzing events with low numbers of particles. Modeling by a two-dimensional Gaussian function can account for circular or elliptical shape only and is therefore mainly suited for the analysis of events involving a high number of particles.

#### Random walk simulation

To accurately mimic the measurement conditions, camera noise as well as temporal blurring were introduced. Camera noise, e. g. read noise and dark noise, follows a normal distribution and can be modeled by fitting a Gaussian function to the baseline images before each stimulation. Temporal blurring was introduced by a 10-fold temporal oversampling and subsequent averaging to account for the motion of particles during the finite exposure times. The standard deviation (8.66 photons) of the modeled camera noise was adjusted by a factor of √10 for 10-fold temporal oversampling. Four particles were generated in the center of the first simulated image (41 x 41 pixels) and Poisson distributed shot noise (mean of 77.73 photons) was added. For the following *n*-1 images, a random 2D walk was simulated using the MatLab random number generator.

### Field emission scanning electron microscopy (FESEM)

Samples were fixed in 2.5% glutaraldehyde (EM grade, Serva) in 0.1 M sodium-cacodylate (Electron Microscopy Sciences) buffer (pH 7.4, adjusted with NaOH) for 2 hours. After washing in sodium-cacodylate buffer, the samples were post-fixed with 1% OsO_4_ (Electron Microscopy Sciences) in sodium-cacodylate buffer for 60 minutes, washed in water, followed by dehydration in a graded series of ethanol (30% up to 100%). Next, the samples were dried via critical point drying using carbon dioxide as transition fluid, followed by 5 nm Pt/C coating by electron beam evaporation using Balzer’s BAF300 (Balzers AG,). Measurements were performed with an S-800 FESEM (Hitachi High-Technologies Corporation).

### Combined light- and dual-beam electron microscopy

Selected samples were used for dual-beam electron microscopy (focused ion beam-scanning electron microscopy / FIB-SEM) after performing live cell imaging of hippocampal neurons transfected with Syp-2xpHl to correlate functionality with ultrastructure. Fiducial markers were introduced with a diamond pen on the back of the glass substrate and with conductive carbon cement (Leit-C, Neubauer Chemikalien) on the front of the glass substrate. For the preparation of samples for FIB-SEM, cells were fixed in 2.5% glutaraldehyde and 2% PFA in 0.1 M sodium cacodylate buffer (pH 7.4, NaOH) for 2 hours. Then, cells were post-fixed with 1% OsO_4_ and 1.5% K_4_[Fe(CN)_6_] (Sigma-Aldrich) in sodium cacodylate buffer for 30 minutes. Afterwards, samples were stained with 1.5% uranyl acetate (Serva) in double-distilled water (overnight at 4 °C), dehydrated in a graded series of ethanol (30% up to 70%), counterstained with 1.5% phosphotungstic acid in 70% ethanol, followed by dehydration in ethanol (70% to 100%). The cell monolayer was soaked with Epoxy resin (Sigma-Aldrich) which was polymerized by incubation for several hours at 40 °C and 60 °C, after draining off any excess resin. The sample was mounted on an aluminum stub using CCC (Conductive Carbon Cement after Göcke, Plano GmbH) and sputter coated with 5 nm platinum using a Balzer MED010 (Balzers AG). The samples were transferred to a Helios 600i FIB-SEM (FEI), and xenapses previously recorded in light microscopy could be relocated with the software Maps 2.1 (FEI). The region of interest was marked with a protective carbon deposition layer and cross-sections were milled perpendicularly with a beam of focused gallium ions at 30 kV, 2.5 nA and imaged with an electron beam at 2 kV, 0.17 nA with an in-lens back-scattered electron detector at a pixel size of 2 nm (via Auto Slice & View software, FEI). Images were aligned in Fiji/ImageJ ^22^ and visualized in Amira (FEI).

### Transmission electron microscopy (TEM) imaging and 3-dimensional visualization

Coverslips were coated with a thin carbon layer (15 to 20 nm) and a 1.5 nm SiO_2_ film on top using Balzer’s BAF300 (Balzers AG). Then, the coat was heat-stabilized at 150 °C for 2 hours. Micropatterning of Nlgn1 and cell culture were performed as described above. On DIV 14-18, cells were either unroofed and fixed at 4 °C as described below, or cells were fixed at RT for 1 hour using 2.5% glutaraldehyde, 4% PFA and 0.01% tannic acid in 0.1 M sodium cacodylate buffer (pH 7.4, adjusted with NaOH). After extensive wash with 0.1 M sodium cacodylate buffer the samples were post-fixed in 1% OsO_4_ with 1.5% K_4_[Fe(CN)_6_] for 1 hour at RT, washed with 0.1 M sodium cacodylate buffer and post-fixed again in 1% OsO_4_ for 1 hour at RT. After washing with double-distilled water, samples were dehydrated in a graded series of ethanol (30% up to 100%). Upon dehydration, the samples were stained *en bloc* with 1.5% uranyl acetate in 70% ethanol for 1 hour. Then, the samples were embedded in Epon 812 epoxy resin (Electron Microscopy Sciences) and blocks were cured for 24 hours at 60 °C. After hardening of the resin, the glass coverslip was removed by heat shock and the exposed surface was further covered with resin and cured for another 24 hours at 60 °C. The areas for ultrathin sectioning were identified with the help of a light microscope, cut out with a diamond saw, trimmed and serially sectioned at 50 - 55 nm with a Leica EM UC7 ultramicrotome (Leica Microsystems GmbH). The ultrathin sections were collected on Formvar-coated slot EM grids and observed with a Philips CM10 TEM (Koninklijke Philips N. V.) operated at 80 kV. Images were acquired with a 16 megapixel bottom-mounted TVIPS F416 CMOS camera controlled by EM Menu software (TVIPS GmbH). Acquired images were then analyzed with MetaMorph software (Molecular Devices). The 3-dimensional reconstructions were done using the Reconstruct freeware program ^23^.

### Scanning electron microscopy (SEM) of unroofed xenapses

Xenapses DIV 16-18, grown on standard 18 mm glass coverslips, were unroofed as described ^24,25^. In brief, cells were washed with calcium-free cold (4 °C) Ringer buffer, then incubated in the same buffer supplemented with 0.1 mg/ml PLL for 15 seconds. Afterwards, the cells underwent a hypo-osmotic shock for 30 seconds in a one-third strength (X3 diluted) internal stabilization buffer (ISB; 70 mM KCl, 30 mM HEPES, 5 mM MgCl_2_; 2 mM EGTA, pH 7.2 adjusted with KOH) ^24^. Subsequently, cells were unroofed in full-strength ISB using a single sonication pulse of about 400 ms duration at the lowest power settings delivered with a 13 mm tip attached to the resonator of Branson 250 Sonifier (Branson Ultrasonics). Immediately after sonication cells were fixed in ISB with 2.5% glutaraldehyde for 15 minutes, thoroughly washed with deionized water, stabilized in 0.1% tannic acid (in deionized water) for another 15 minutes and, after additional washing, post-fixed with aqueous uranyl acetate 0.1% for 15 minutes. After washing, cells were dehydrated in an ethanol series and critically-post dried from 100% ethanol using carbon dioxide as transition fluid. After drying, regions containing unroofed xenapses were identified using phase-contrast light microscopy and small pieces of coverslips were cut out with a diamond pen. These samples were glued to aluminum sample carriers for SEM using CCC (Conductive Carbon Cement after Göcke, Plano GmbH) and subsequently coated with Pt/C 2.1 nm by electron beam evaporation using Balzer’s BAF300 (Balzers AG). SEM was performed with a Hitachi S-5000 FESEM (Hitachi High-Technologies Corporation) at 30 kV using a secondary electron detector. Images were acquired and digitized using DISS5 (Point Electronic).

### Freeze-fracture of xenapses

Xenapses DIV 16-18, grown on 12 mm glass coverslips, were washed in standard Ringer buffer. Then, cells were shortly dipped in double-distilled water and placed cell-side down on 10 mm custom-made SEM copper stubs coated with a layer of cold water fish skin gelatine (20% in water, Sigma-Aldrich). Cells were immediately frozen by plunging into liquid nitrogen (LN2). The frozen samples were transferred from the LN2 into the pre-cooled loading station EM VCM, mounted on the sample holder for SEM stubs and, using vacuum cryo-transfer EM VCT500, were introduced into the EM ACE900 (all Leica Microsystems) for freeze fracturing. The freeze fracture was performed at −120 °C in manual mode by lifting off the coverslip with a knife. The sample was freeze dried at −80 °C for 5 min, coated by unidirectional evaporation at 45° with Pt/C 2 nm, and baked out with carbon 5 nm at 90°. The Pt/C replica was released onto the surface of double-distilled water at RT and picked up on a drop with a stainless-steel loop and transferred onto Formvar-coated EM grids (mesh 75-100) for TEM observations.

---

**Figure S1.**
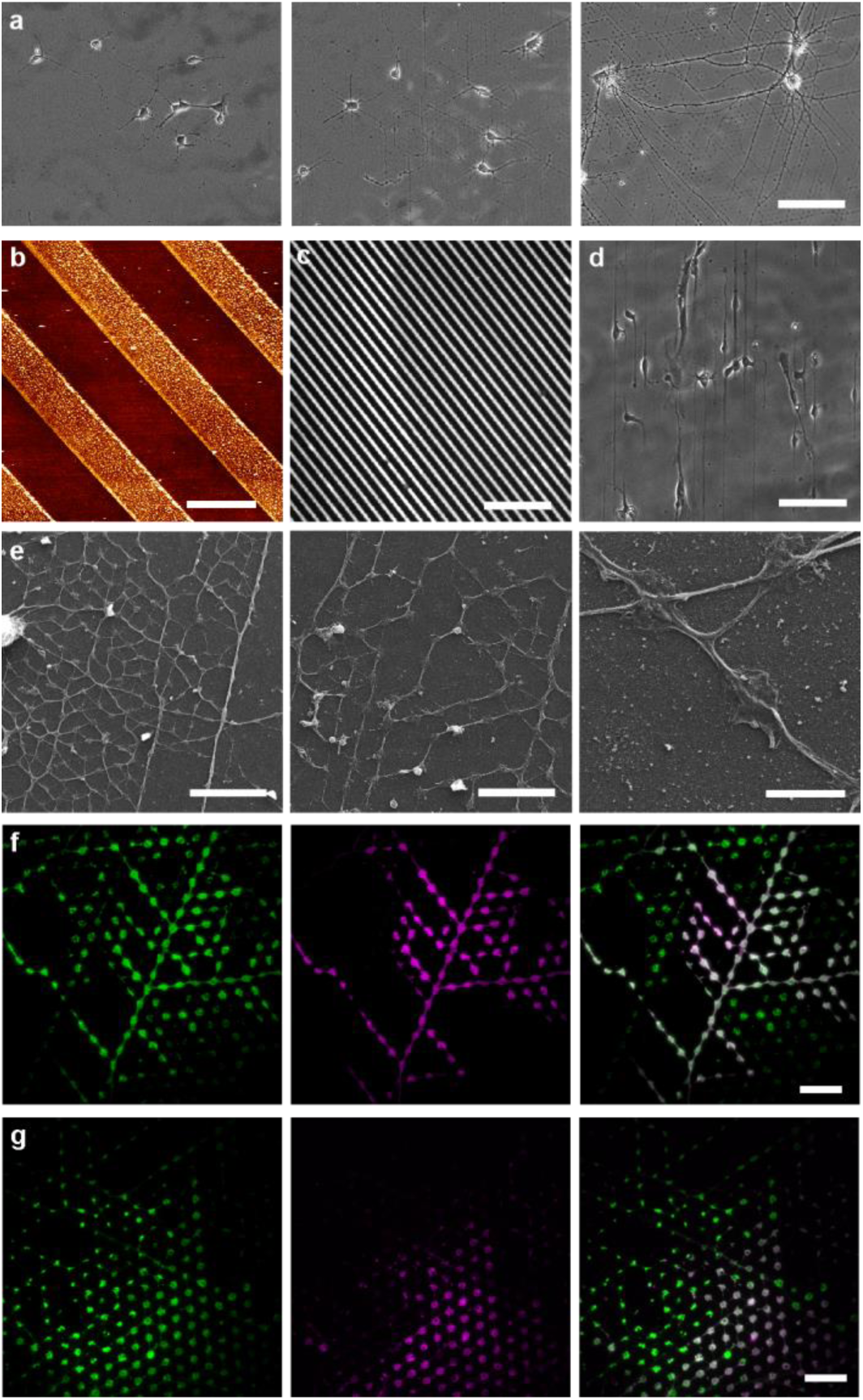
Microstructured coverslips functionalized with Nlgn 1 serve as artificial host substrates to induce presynapses. **a**, Phase contrast images of hippocampal neurons growing on functionalized substrates after two (left), three (centre) and eight days (right) in culture (DIV), scale bar 100 μm. **b-d**, Visualization of the stripe pattern used for the functionalization of coverslips. **b**, Microstructured substrate imaged via AFM (scale bar 10 µm), **c**, Labelling of the functionalized substrate with Nlgn1 antibody, scale bar 100 μm and **d**, Phase contrast images of hippocampal neurons (2 DIV) growing on the stripe pattern. **e,** Scanning electron microscopy images of hippocampal neurons (16 DIV) growing on the substrate functionalized with small stamps (2,5 µm dot diameter, 3 µm distance between dots) with zoom-in of single axons forming xenapses (right). Scale bars 20 µm, 12 µm and 3 µm, respectively. **f and g**, Hippocampal neurons (8 DIV) labelled with antibodies against Syp1 (green) and against VGAT (magenta, **f**) or VGlut (magenta, **g**) indicate the presence of both GABAergic as well as glutamatergic synapses. Scale bars 20 μm.

**Figure S2.**
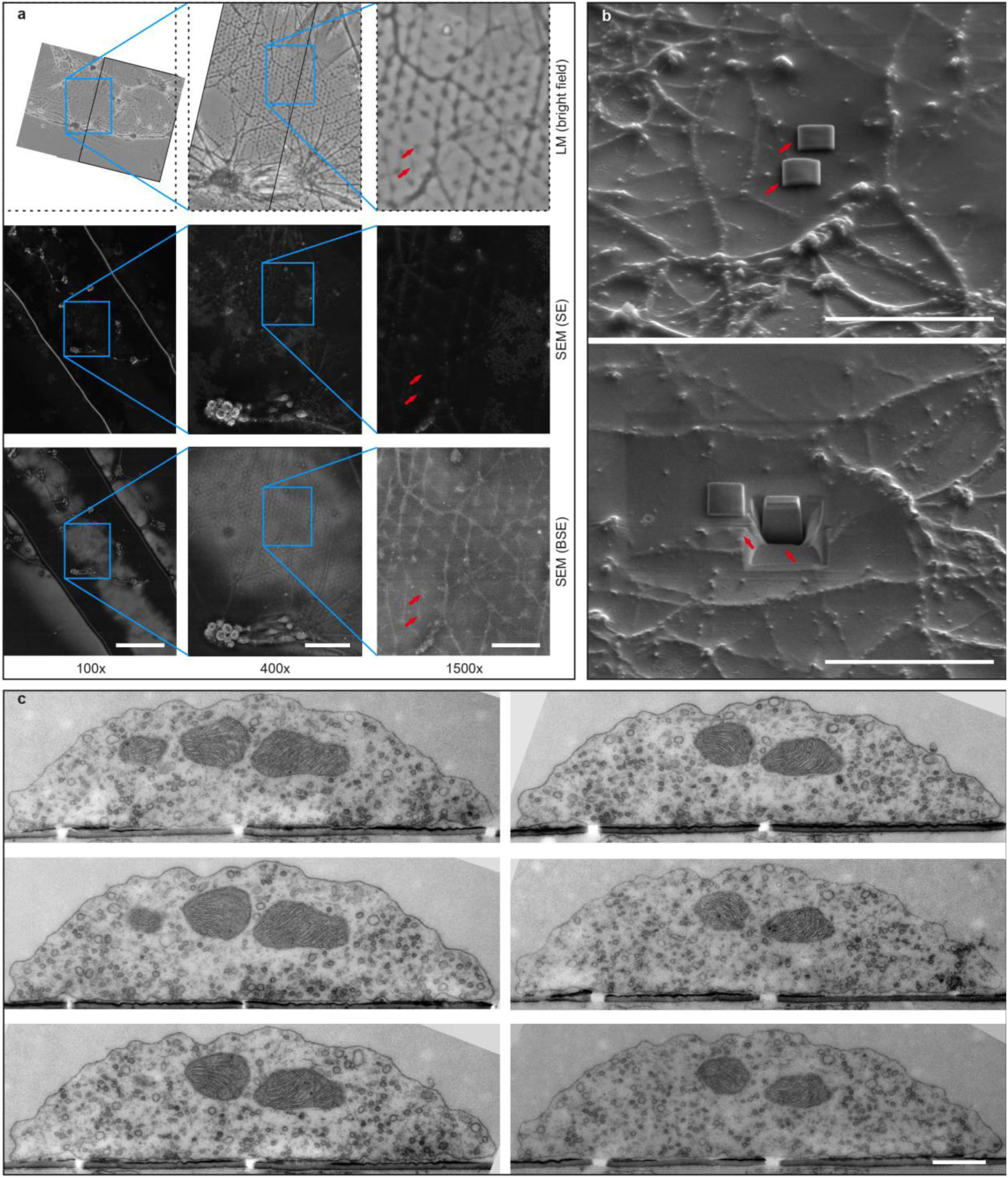
Xenapses form directly on the functionalized substrate with many vesicles docked at the bottom facing the Nlgn1 protein pattern. **a**, Re-localization of specific xenapses previously imaged live with fluorescence microscopy for FIB-SEM measurements. Xenapses are imaged by light microscopy (LM, bright field illumination) at multiple magnifications, fixed and embedded (top). This enables tracing of the respective areas in electron microscopy (EM, with both secondary electrons (SE, middle) and back-scattered electrons (BSE, bottom)). Arrows indicate the xenapses of interest. Scale bars 300 µm, 70 µm and 20 µm, respectively. **b**, Xenapses are prepared for FIB-SEM imaging. Arrows point to the xenapses of interest, covered by a protective layer of carbon (top) and milled with a focused beam of gallium ions (bottom). Scale bar 20 µm. **c**, Exemplary TEM serial sections used for 3D reconstruction of a single xenapse (figure 2b). Scale bar 500 nm.

**Figure S4.**
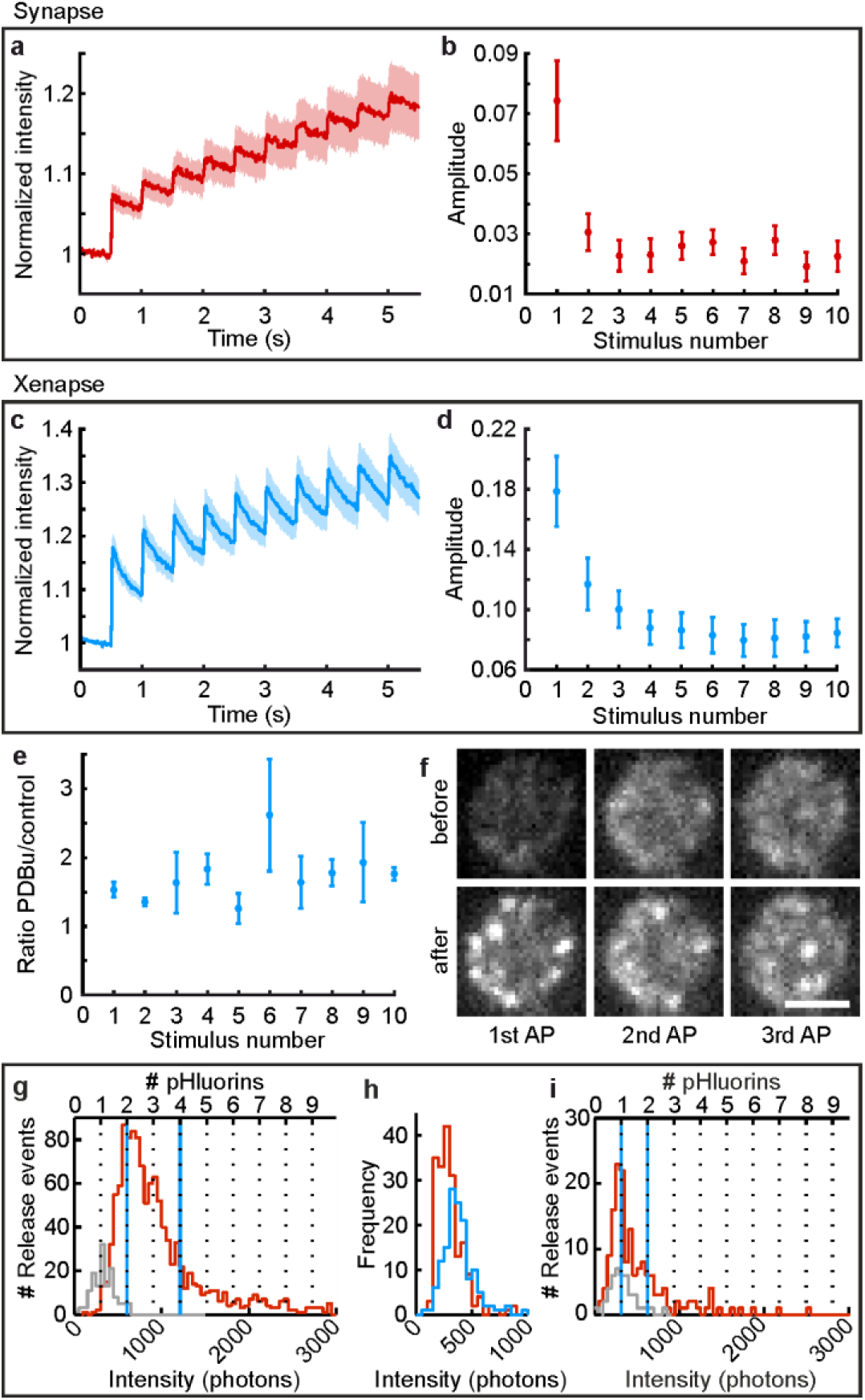
Release characteristics of Syp1-2xpHl-transfected xenapses to trains of and single APs. **a to d**, Average Syp-2xpHl responses to 10 APs (2 Hz) in, **a**, synapses (n = 8 regions each comprising more than 100 boutons, 4 biological replicates) and in, **c**, xenapses (n = 10 regions each comprising more than 20 boutons, 4 biological replicates). The increase in amplitude for each AP in synapses (**b**) and xenapses (**d**) was determined by averaging the intensities of two frames post-stimulus and subtracting the intensity before the stimulus. All errors in s.e.m. **e**, Ratio of responses of xenapses after and before application of PDBu (n = 3 regions, 3 biological replicates). **f**, Representative xenaptic bouton (17 DIV) expressing Syp1-2xpHl before (top row) and after (lower row) stimulation with single APs. Scale bar 1 µm. **g**, Accumulated intensity histogram for 4448 single localized fusion events (red) and baseline blinking events (gray) of Syp1-2xpHl expressing xenapses (13 regions, 10 biological replicates). Syp1-2xpH contains two pH per molecule, thus events with equal or less than two pHl are considered to represent fusion of SVs with only one Syp1-2xpHl molecule, while events with more than four pHl are referred to as fusion of SVs harboring multiple Syp1-2xpHl molecules. The respective borders are shown in blue. **h**, Fluorescence intensity histograms of single pHl (red) and EGFP (blue) proteins immobilized on coverslips. The average fluorescence of single pHl molecules is nearly identical to that of baseline blinks in xenaptic recordings, and is used to convert fluorescence intensities in **g** to numbers of pHl molecules (upper abscissa). **i,** Accumulated intensity histogram of evoked fusion events (red) and baseline blinking events (gray) of Syt1-pHl expressing xenapses (17 DIV). Syt1-pHl contains one copy of pHl, thus events with more than two pHl are considered as fusion of SVs harboring multiple Syt1-pHl molecules (borders for intensities of single and multiple pHl events marked by blue lines).

**Figure S5.1.**
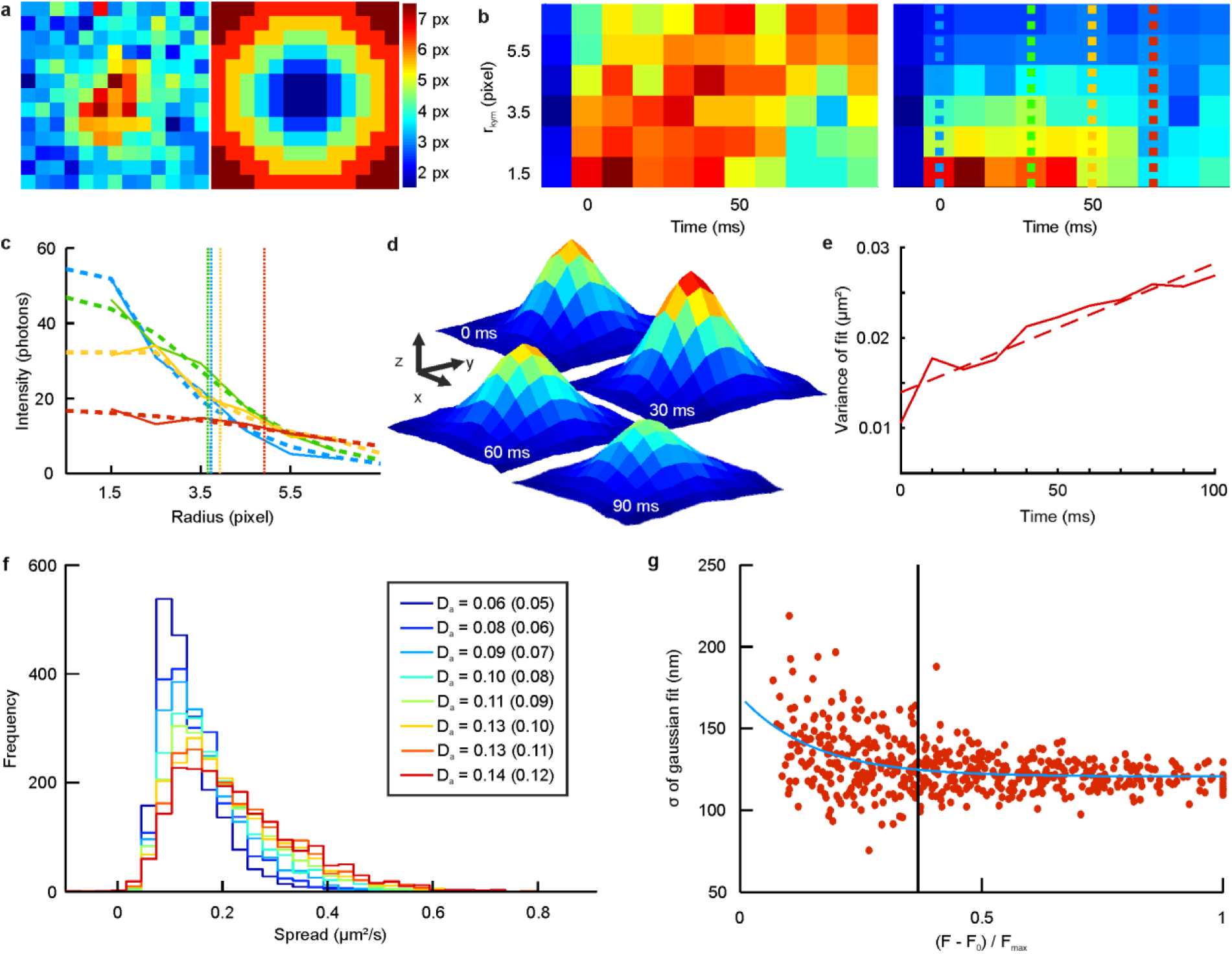
Recordings of single vesicles containing multiple Syp1-pH reveal diffusional dispersion of SV proteins post fusion. **a**, Generation of radial kymographs. Pixel values from an exemplary fusion event (left) are extracted using ring-shaped masks (right) and summed up. Each ring correlates with the distance from the center of the region of interest (color code, right). **b**, Radial kymograph (left) from summed pixel values for all ring-shaped masks (ordinate) and every time step. Weighted kymograph (right) from dividing the summed pixel values by the number of pixels in each ring. Vertical dashed lines refer to distinct time points (blue - 0 ms, green - 30 ms, yellow - 50 ms or red – 70 ms) after stimulation. **c**, For selected time points of the weighted kymograph (i.e. along the vertical lines in **b**, right panel), pixel intensities are fit with a logistic function to yield the width at half maximum. Pixel intensities extracted from the weighted kymograph are displayed as solid lines colored according to vertical lines in **b**, the logistic fits as dashed lines. The width at half maximum is marked by dotted lines. **d and e**, Spread of superimposed events at different time points to determine apparent diffusion constants. **d**, Average signal of events with single Syp1-2xpHl for different time points. Scale bars 200 nm (x and y) and 0.2 a.u. (z). **e**, Widths (variance) of Gaussian fits to averaged events in **d** for the first 10 frames (100 ms, solid line). A linear fit (dashed line) yields an apparent diffusion coefficient of 0.0716 µm^2^/s for events with single Syp1-2xpHl. **f**, Relationship between spread parameter from radial kymograph analysis and diffusion coefficient. Plotted are histograms of spread values from radial kymograph analyses of individual Monte Carlo simulations (random walks) for four simultaneously released molecules with given diffusion constants. Values in the legend give the median spread of the particular histogram with the input diffusion constants (for the simulations) in brackets (in µm^2^/s). **g**, Widths of Gaussian fits to images of beads (40 nm) diffusing in and out of the evanescent field versus normalized intensity. Widening of the PSF due to vertical movement out of the evanescent field (exponential fit, blue) is negligible, as the PSFs standard deviation increases by less than 4 % (estimated from exponential fit to data for an intensity drop to I_max_/e (vertical line)). Imaged fluorophores fade away before broadening of the PSF becomes visible. Thus, the observed spread of fusion events cannot be explained by movement out of the evanescent field, but reflects lateral dispersion.

**Figure S5.2.**
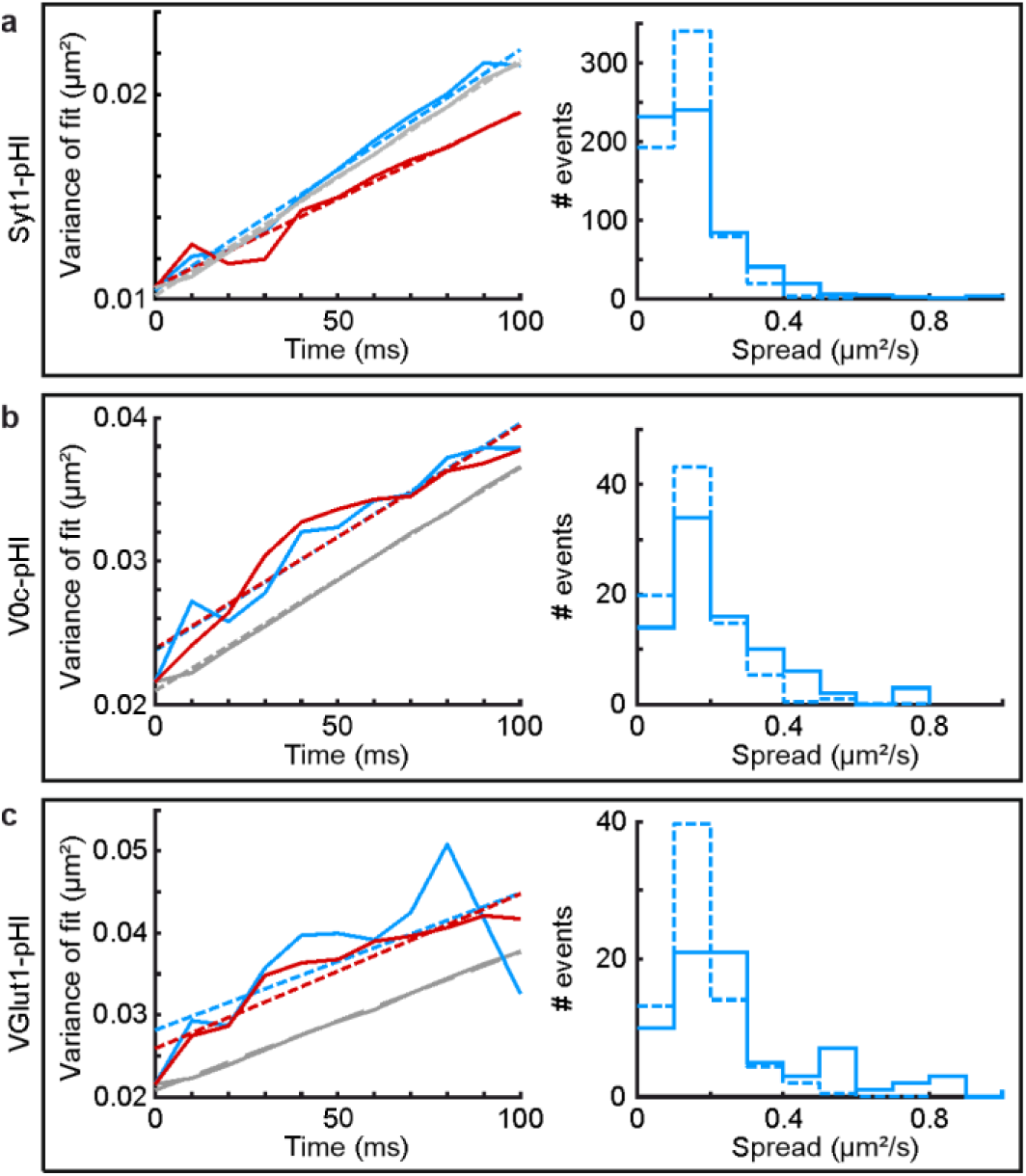
Recordings of single vesicles containing multiple copies of Syt1-pHl, V0c-pHl (V-ATPase) or VGlut1-pHl reveal diffusional dispersion after fusion. **a**, Diffusion analysis of evoked release events in xenapses expressing Syt1-pHl. Left: widths (variance) of Gaussian fits to centered and averaged events with multiple copies (solid blue) and single copies (solid red) for the first 10 frames (100 ms). Linear fits yield diffusion coefficients of 0.0586 µm^2^/s for events with multiple Syt1-pHl copies (dashed blue) and of 0.0426 µm^2^/s for single copy events (dashed red). Widths of Gaussian fits to the average of 1000 simulated events using the measured diffusion coefficient of 0.0586 µm^2^/s (grey solid) can be fit with a line (grey dashed), yielding a diffusion constant of 0.0574 µm^2^/s. Right: histograms of spread values of 639 multiple copy events from radial kymograph analysis (solid) and of 1000 simulated events (dashed). The median spread values are 0.0799 µm^2^/s for measured and 0.0809 µm^2^/s for simulated events. Note that the histogram of simulated data was normalized to the integral of the histogram of measured data (n = 31 regions, 3 biological replicates). **b**, Diffusion analysis of evoked release events as in **a** of xenapses expressing V0c-pHl. Left: Linear fits yield diffusion coefficients of 0.0793 µm^2^/s for events with multiple copies (dashed blue), of 0.0778 µm^2^/s for single copy events (dashed red), and of 0.0778 µm^2^/s (dashed gray) for simulated multiple-copy events. Right: histograms of spread values of 85 multiple copy events (solid) and of 1000 simulated events (dashed). The median spread values are 0.1258 µm^2^/s for measured and 0.0879 µm^2^/s for simulated events (n = 11 regions, 3 biological replicates). **c**, Diffusion analysis of evoked release events as in **a** of xenapses expressing VGlut1-pHl. Left: Linear fits yield diffusion coefficients of 0.0836 µm^2^/s for events with multiple copies (dashed blue), of 0.0943 µm^2^/s for single copy events (dashed red) and of 0.0840 µm^2^/s (dashed gray) for simulated multiple-copy events. Right: histograms of spread values of 74 multiple copy events (solid) and of 1000 simulated events (dashed). The median spread values are 0.1652 µm^2^/s for measured and 0.0969 µm^2^/s for simulated events (n = 10 regions, 9 biological replicates).

**Figure S5.3.**
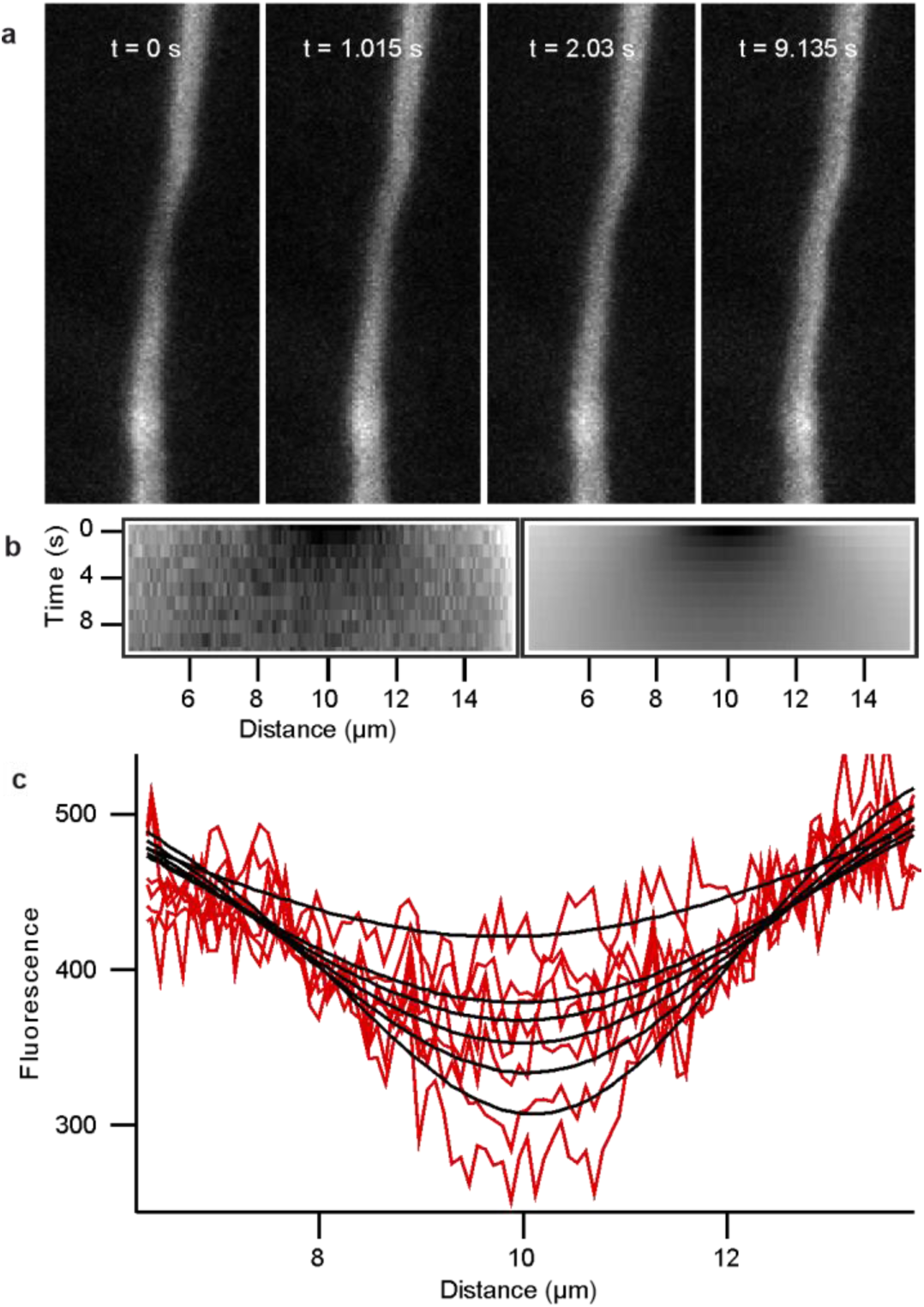
FRAP experiments in the axon yield a diffusion constant of 0.38 µm^2^/s for Syp1-pH. **a**, Time lapse series of fluorescence recovery in an axonal branch after bleaching by line scans perpendicular to the axon (bleaching at 2 mW at the objective back pupil). **b**, Kymograph of the axon depicted in **a** showing the fluorescence recovery post bleach (left). Right: best least-square fit to the measured kymograph for one-dimensional diffusion. The evolution of concentration C in space and time in the kymograph was fit by the equation (Crank, J. *The Mathematics of Diffusion* (Oxford University Press, Oxford, 1975), p. 63, eq. (4.56))

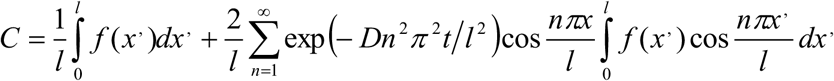

using the fluorescence profile along the axon measured in the first image post bleach as initial distribution f(x) and assuming impermeable boundaries at both ends of the axon (top and bottom of images in **a**. Least square fitting with a Levenberg-Marquardt routine yielded a diffusion constant of 0.52 µm^2^/s. **c**, Exemplary spatial profiles and corresponding fits (D = 0.52 µm^2^/s) from **b** for t = 0, 1.015, 2.03, 3.045, 4.06, 5.075, and 10.15 s. Five additional experiments yielded an average diffusion constant of 0.38 ± 0.10 µm^2^/s.

